# Origin and Correlates of Viral Rebound in SIV-Infected Rhesus Macaques Following ART Discontinuation

**DOI:** 10.1101/2025.08.30.673277

**Authors:** Irena V. King, Malika Aid, Emek Kose, Taina T. Immonen, Charles A. Goodman, Christine M. Fennessey, Alessandro Colarusso, Victoria E. K. Walker-Sperling, Erica N. Borducchi, Romas Geleziunas, William J. Rinaldi, Melissa J. Ferguson, Louis J. Picker, Jeffrey D. Lifson, Brandon F. Keele, Dan H. Barouch

**Author notes:** Corresponding author: D.H.B.

## Abstract

The vast majority of persons living with HIV-1 who discontinue antiretroviral therapy (ART) demonstrate viral rebound, but the tissue-level events that lead to rebound viremia are poorly understood. Here we report the origin, dynamics, and correlates of viral rebound in 16 rhesus macaques (RMs) infected with molecularly barcoded SIVmac239M, treated with ART for 70 weeks, and necropsied on day 12 after ART discontinuation. Barcode analysis of plasma following ART discontinuation identified 1 to 38 rebounding barcode-defined viral lineages per animal, with 1 to 4 rebounding lineages contributing to first measurable rebound viremia. Analysis of barcode viral RNA (vRNA) expression in necropsy tissues revealed presumptive anatomic origin sites for 56 of 175 total rebounding viral lineages, with significant enrichment in the gastrointestinal (GI) tract and GI-associated lymph nodes. Daily transcriptomic and proteomic profiling in peripheral blood following ART discontinuation showed upregulation of pathways related to T cell signaling, cytokine responses, and cellular metabolism prior to detectable rebound viremia. These data suggest that viral rebound following ART discontinuation is initiated by local tissue replication of a limited number of clonal lineages, followed by systemic expansion of the initial rebounding lineages and serial initiation of replication of multiple additional clonal lineages. These findings provide mechanistic insights into the processes that result in viral rebound following ART discontinuation and will contribute to next generation HIV-1 cure strategies.

---

The replication-competent viral reservoir (RCVR) that persists despite extended ART and can give rise to viral recrudescence following ART discontinuation remains the key obstacle to achieving an HIV-1 cure. The RCVR includes latently infected CD4+ T cells distributed across diverse tissues and is established early during infection and is not eliminated by ART, enabling viral recrudescence following ART discontinuation in the vast majority of individuals.^2–5^ Current methods for quantitating virus that persists on ART have limited capacity to predict viral rebound due to the presence of defective proviruses, incomplete latency, and other factors.^1,5^ Using molecularly barcoded SIVmac239M,^6,7^ we investigate viral rebound dynamics in RMs following discontinuation of ART that was initiated during acute infection.^6^ We define the timing and number of barcode lineages that contribute to viral rebound following ART discontinuation and identify presumptive anatomic origin sites for a subset of the rebounding clonal lineages. Parallel transcriptomic and proteomic analyses of peripheral blood following ART discontinuation identified early host inflammatory and metabolic pathways associated with viral rebound.

## Study design

18 outbred, Indian-origin adult male and female RMs (*Macaca mulatta*) (*n*=6 per group) were intravenously infected with 5,000 infectious units (IUs) of molecularly barcoded SIVmac239M. To achieve varying levels of RCVR seeding, ART was initiated on day 6, 9, or 12 following infection. The ART regimen consisted of daily subcutaneous injections of tenofovir disoproxil fumarate (TDF; 5.1 mg/kg/day), emtricitabine (FTC; 40 mg/kg/day), and dolutegravir (DTG; 2.5 mg/kg/day) pre-formulated in a 15% (v/v) kleptose solution at pH 4.2.^8^ Animals were treated with daily ART for 70 weeks with longitudinal plasma viral loads and peripheral blood mononuclear cell (PBMC) viral RNA (vRNA) and viral DNA (vDNA) measurements. Following ART discontinuation at week 70, animals were monitored daily for 12 days and then underwent comprehensive necropsy. A median of 62 tissue samples per animal, including GI, lymphoid, and non-lymphoid sites, were harvested at necropsy to assess total vDNA and vRNA levels and clonotypic barcode representation in vDNA and vRNA.

## Virologic and immunologic dynamics on ART

Following ART initiation, plasma viremia was suppressed to <15 copies/mL at most timepoints after day 120 for animals in the day 6 and day 9 ART initiation groups (**Fig. 1A, B**). Time to achieve viral suppression was longer and more variable in animals that initiated ART on day 12, consistent with higher plasma viral loads prior to ART initiation (**Fig. 1C-E**). The day 12 group also exhibited more frequent on-ART blips. Two RMs, one each from the day 6 (DHTG) and day 9 (DHPG) groups were euthanized for procedure-related complications during the ART treatment period. vDNA and vRNA in PBMCs declined during ART, with vRNA declining more rapidly than vDNA, as expected (**Fig. 1F-G**). Intact proviral DNA in PBMC, detected using an SIV intact proviral DNA assay (IPDA), was lower in RMs that initiated ART on day 6 (**Fig. S1**).^4,9^ On-ART biopsy tissue analyses revealed early and widespread detection of vDNA and vRNA in all groups across lymphoid and GI tissues (**Fig. 1H–I**; **Fig. S2**).

**Figure 1.**
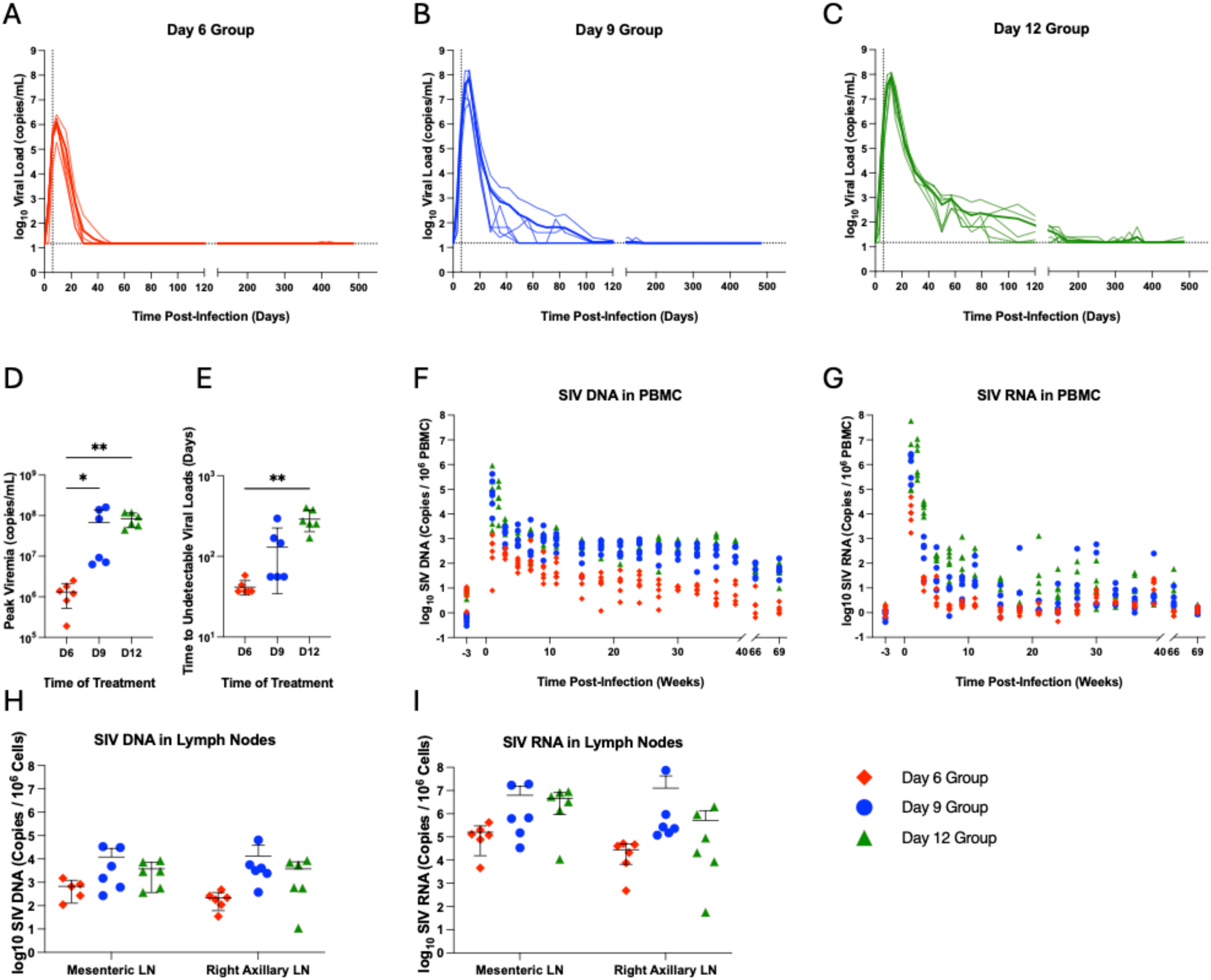
Plasma viral loads and cell-associated SIV DNA and RNA dynamics following ART initiation in SIVmac239M-infected animals. Plasma viral loads are shown in individual animals infected with SIVmac239M and treated with ART beginning on (A) day 6 (*n*=6; red), (B) day 9 (*n*=6; blue), or (C) day 12 (*n*=6; green). Vertical dashed lined is the time of ART initiation and horizontal dashed line is the limit of detection of 15 copies/mL. (D) Peak viral loads and (E) time to plasma viral loads <15 copies/mL. The levels of (F) cell-associated viral DNA (vDNA) and (G) cell-associated viral RNA (vRNA) in individual animals were measured in PBMC. Levels of (h) vDNA and (i) vRNA in mesenteric and right axillary lymph node biopsies three days after ART initiation in each respective group. Black lines indicate mean ± standard deviation. * *p*<0.0332, ** *p*<0.0021.

We next assessed SIV-specific antibody and T cell responses during ART suppression. Earlier ART initiation generally resulted in lower SIV Env-specific antibody titers (**Fig. S3A-D**). SIV-specific cellular immune responses were evaluated using IFN-γ ELISPOT and intracellular cytokine staining assays. SIV-specific T cell responses were undetectable throughout follow-up in the day 6 ART initiation group and in most day 9 and day 12 ART initiation RM at week 33 post-infection (**Fig. S3E**), but Gag-specific responses were observed at week 66 in a subset of day 9 and day 12 ART initiation RM (**Fig. S3F-G**). At necropsy, Gag-specific IFN-γ^+^ and TNF-α^+^ CD4 T cell responses were detected in the spleen in most animals (**Fig. S3H-I**). These findings suggest that later ART initiation may result in greater humoral and cellular immunity, potentially due to increased antigen exposure.^10^

## Viral rebound after ART discontinuation

At week 70, ART was discontinued in all 16 RMs, and plasma viral loads were monitored on days 0, 2, 4, and then daily until day 12 following ART discontinuation to assess viral rebound dynamics. Two animals (TP5, DHHN) did not rebound by day 12 (**Fig. 2A**). Among the 14 animals that showed plasma viremia, time to viremia >50 copies/mL was observed between 6 and 12 days after ART discontinuation (**Fig. 2A; Fig. S4**). Rebound viremia at necropsy was <10^3^ copies/mL in 3 of the 14 viremic RM and was between 10^3^ and 10^7^ copies/mL in the remaining 11 RMs. Calculated plasma viral load rebound growth rates varied for individual RMs (mean 1.75, range 1.10-2.65 log RNA/day), with no significant differences between groups (Kruskal-Wallis; *p*=0.22; **Table S1**).

**Figure 2.**
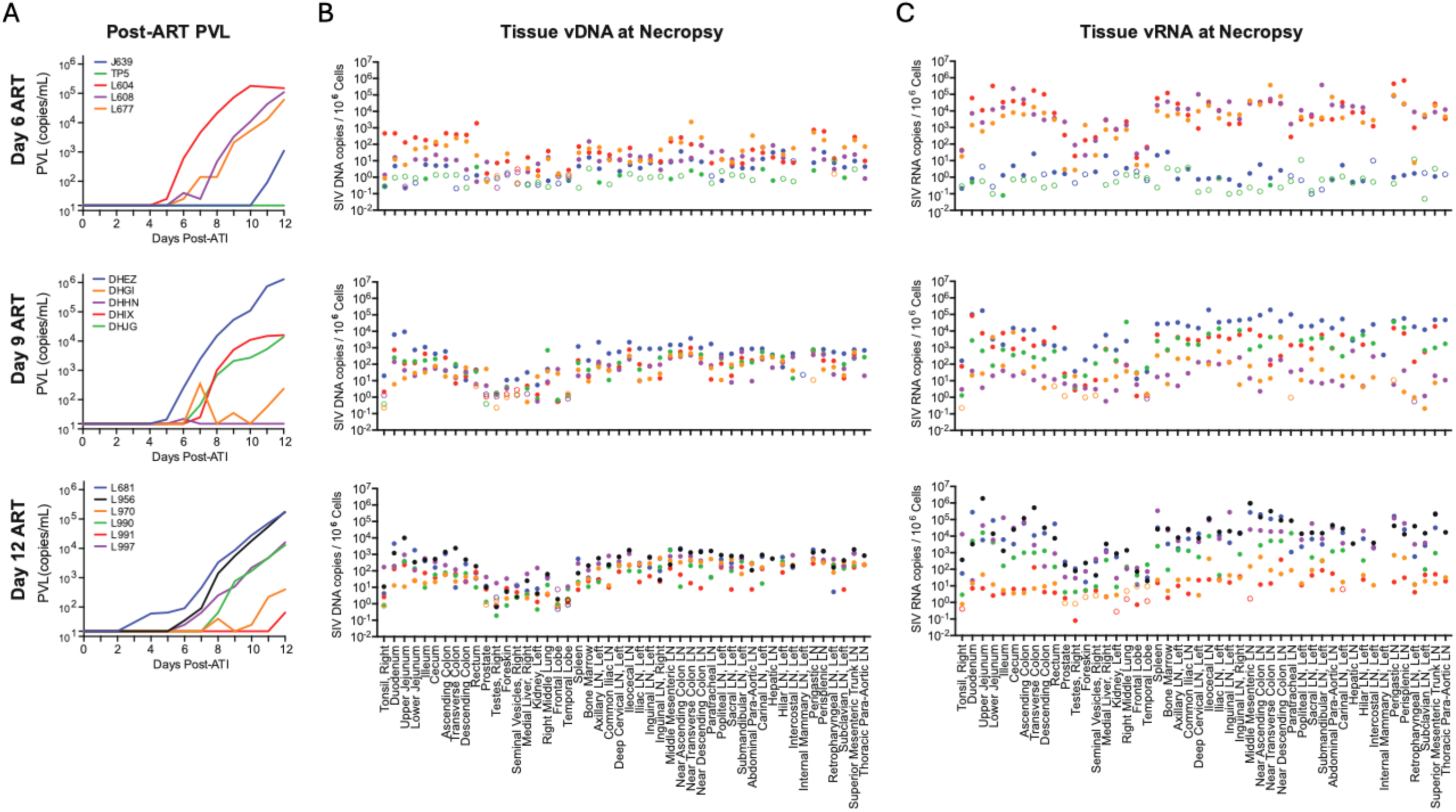
Plasma viral loads and necropsy tissue vRNA and vDNA levels following ART discontinuation. (A) Plasma viral loads for 12 days following ART discontinuation at week 70 are shown for the day 6 group (*n*=5; 4/5 rebounders), day 9 group (*n*=5; 4/5 rebounders), and day 12 group (*n*=6; 6/6 rebounders). The limit of detection is 15 SIV RNA copies/mL. Tissue distribution of vDNA (B) or vRNA (C) at necropsy is shown as the number of viral copies per million cells. Unfilled circles represent samples in which no replicates were positive for vDNA or vRNA, and an imputed threshold value was plotted to indicate levels below the detection limit.

Sequencing of rebound plasma virus showed 1 to 38 barcode lineages per animal (**Fig. S4; Table S1**) with 1 to 4 lineages (mean 2.4) contributing to the initial measurable rebound viremia. Individual RM growth curves and barcode proportions were used to calculate the average reactivation rate per animal (**Table S1**).^6,11^ The overall rate for all animals was 1.4 (range 0.12-5.67) reactivation events/day that led to detectable rebound viremia, with no significant differences between groups (Kruskal-Wallis, *p*=0.34).

Total vDNA and vRNA quantitation was performed for tissue samples obtained at necropsy on day 12 following ART discontinuation. A standardized tissue collection protocol was used to obtain a median of 62 (range 51-85) independent tissue samples from each animal, including GI tissues, GI-associated and other lymphoid tissues, as well as non-GI/non-lymphoid tissues (**Table S2**). Viral quantitation revealed vDNA (**Fig. 2B**) and vRNA (**Fig. 2C**) in multiple tissues, including GI tract and lymphoid tissues, with lower levels of vDNA in non-GI/non-lymphoid tissues, such as reproductive organs, lungs, liver, and brain.

Barcode sequencing of vDNA and vRNA was employed to assess viral lineages replicating within individual necropsy tissue specimens, with barcode sequences obtained from a mean of 39 (range 11-45) samples per animal, making it possible to demonstrate some individual barcode lineages identified in primary infection plasma viremia prior to ART initiation as contributing to viral rebound in plasma and also present in tissues following ART discontinuation. Consistent with previous studies and a companion study,^6,11,12^ the representation of individual barcode lineages in plasma viremia during primary infection was predictive of the probability of those barcode lineages being represented in rebound viremia (**Fig. S5**; **Table S3A**; *p*<0.001, 2-sided Wald test). To assess if rebounding barcode lineages showed evidence of viral replication in tissues compared to barcode lineages not identified in rebound viremia, we compared the cumulative vRNA and vDNA levels of each individual barcode representation in all necropsy tissues to the contributions of individual barcodes in primary peak viral load, with rebounding lineages (heat map color) readily distinguishable from non-rebounding lineages (grey) (**Fig. S6**). Barcode lineages found in rebound plasma viremia had significantly higher total tissue vRNA and vDNA levels compared to non-rebounding lineages (log_10_ vRNA median 4.9 vs. 1.9; vDNA median 3.6 vs. 1.7; *p*<2x10^-16^, Wilcoxon rank-sum test). Furthermore, total vRNA and vDNA levels in tissues for individual barcodes predicted rebound viremia levels for those barcodes (**Table S3B**; *p*<0.001, *p*<0.001, 2-sided Wald test), providing evidence that individual barcode lineages actively replicating in tissues following ART discontinuation are contributing to rebound viremia.

## Viral barcode clonotype dynamics in tissues

The use of a barcoded virus enables the possibility of not only tracking the contributions of individual barcode clonotypes to rebound viremia, but also potential identification of the tissue sites of origin of such rebounding barcodes. In a companion study,^12^ RMs necropsied on ART were used to define the limits of barcode-level vRNA expression on ART and to establish a 99% predictive interval for vRNA expression in the absence of viral replication. Analysis of tissues from RM necropsied 5 or 7 days after ART discontinuation revealed rare tissues with barcode vRNA exceeding this 99% prediction interval as sites of initial viral replication after ART discontinuation and when those barcodes matched barcodes present in the earliest stages of rebound viremia (<30 copies/mL), the tissue origin site for these lineages could be identified.

In the present study, necropsies were performed 12 days after ART discontinuation with rebound plasma viremia between 10^3^ and 10^7^ copies/mL in 11 of 14 RM that were viremic at necropsy, often with vRNA expression of barcode lineages present in rebound plasma viremia identified in multiple tissues. To try to determine the tissue origin sites for rebounding barcode lineages in these conditions, an alternative method was employed using a machine-learning clustering approach to identify outlier tissue sites where vRNA expression compared to other tissues also expressing the same rebounding barcode vRNA exceeded the level defining disseminated replication (determined from non-outlier tissues of all rebounding barcodes), thereby implicating those individual tissue specimens as presumptive tissue origin sites (**Figs. 3-5**, **Figs. S7-17**).

**Figure 3.**
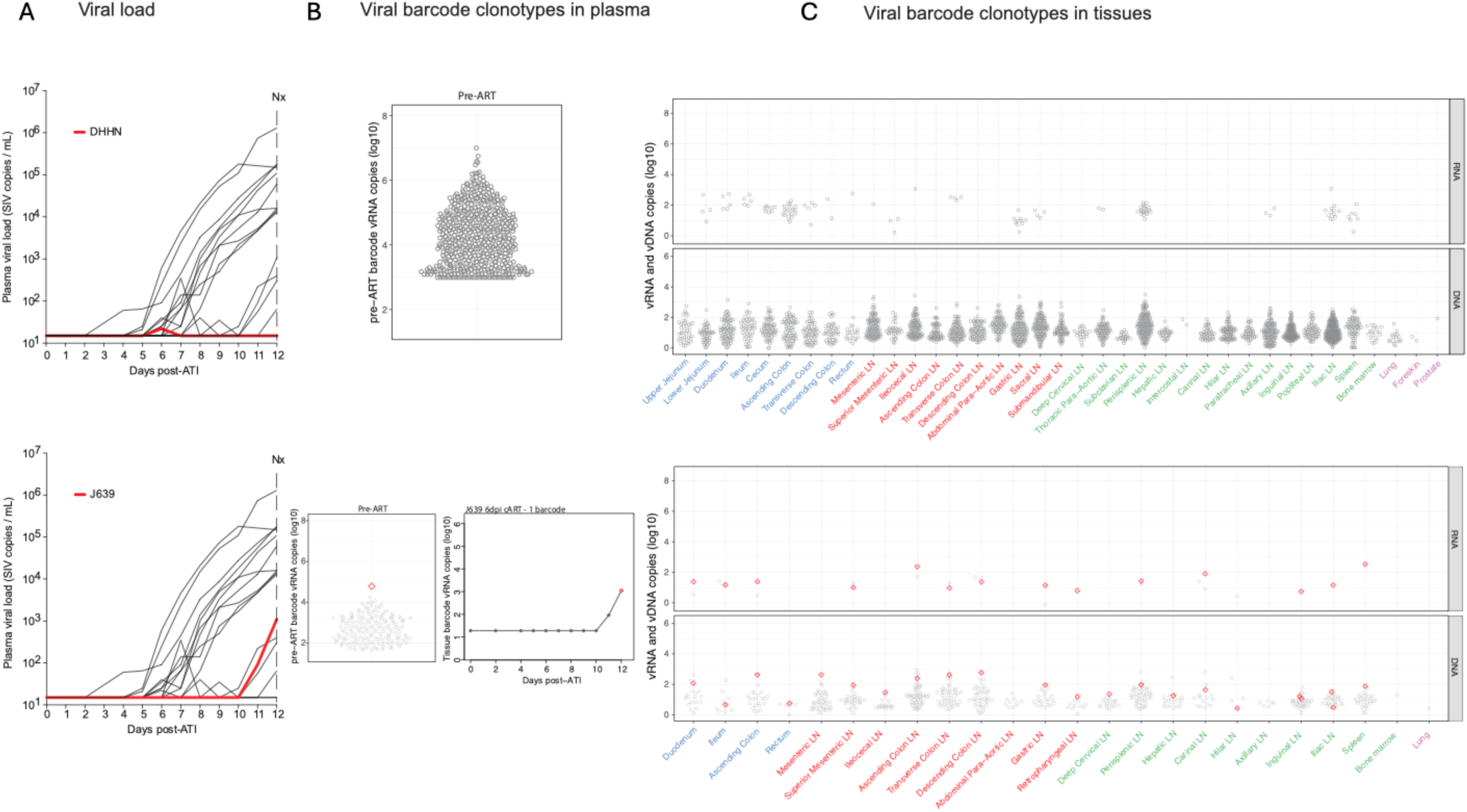
Viral barcode clonotypes in necropsy tissues from a non-rebounder RM DHHN and a single-barcode rebounder RM J639. (A) Plasma viral load dynamics following ART discontinuation for RMs DHHN (top) and J639 (bottom); red lines indicate the viral load trajectories for RM DHHN and RM J639. RM DHHN did not show viral rebound during the 12 day monitoring period. (B) Proportional distributions of viral barcode clonotypes in plasma viremia during primary infection prior to ART initiation for RMs DHHN (top) and J639 (bottom); for J639, the barcode detected in rebound plasma (bottom left panel) is highlighted in red. The bottom right panel shows calculated rebound viral growth curves for the rebounding barcode lineage with estimated time to a single copy in rebound viremia indicated. (C) Necropsy tissue distribution of vRNA and vDNA barcode clonotypes from primary infection for each animal. For RM J639 (bottom), the plasma rebound barcode (BC.2497) was detected in gut and lymphoid tissues, although no origin site was identified. Grouped tissue categories are: GI tract (blue), GI tract draining lymph nodes (red), non-GI lymph tissues (green), non-lymphoid tissue (purple).

Two animals (DHHN, TP5) did not exhibit plasma viral rebound and also did not have sufficient vRNA levels in tissues on day 12 following ART discontinuation for outlier analysis, likely due to early ART initiation (**Fig. 3**, top; **Fig. S7**, top). Two animals (J639, L970) showed initial plasma viral rebound on day 11 following ART discontinuation and demonstrated a single barcode vRNA in plasma at necropsy (**Fig. 3**, bottom; **Fig. S7**, bottom). Animal J639 showed the same barcode (BC.2497; red) in primary viremia prior to ART initiation, in rebound plasma viremia on day 12 following ART discontinuation, and in GI and lymphoid tissues at necropsy, but no tissue origin site was identified (**Fig. 3B-C**, bottom). Animal L991 showed initial plasma viral rebound on day 12 following ART discontinuation with two barcodes (BC.897, BC.2604) that were identified in peak primary infection plasma and also detected in rebound plasma viremia (**Fig. S8**). Tissue analysis detected vRNA and vDNA for both of these barcodes in multiple tissues.

Animal L604 showed measurable plasma viremia on day 6 following ART discontinuation (**Fig. 4A**). Three barcodes were identified in rebound viremia with a level of 180,000 copies/mL at necropsy (**Fig. 4B**). Barcode analysis of primary infection showed that these three barcodes were relatively abundant in primary viremia prior to ART initiation (**Fig. 4B**), although the order of detectable barcode emergence in plasma following ART discontinuation did not correspond to their abundances in plasma prior to ART initiation. Necropsy tissue analysis revealed that the first barcode detectable in rebound viremia (red, BC.15) was already widely disseminated, whereas the second (blue, BC.4095) and third (purple, BC.1192) barcodes identified in rebound viremia were more localized to the lower jejunum and perisplenic lymph node (LN), respectively (**Fig. 4C**), which were identified as tissue origin sites for the second (blue, BC.4095) and third (purple, BC.1192) detectable rebounding lineages, respectively (**Fig. 4D**).

**Figure 4.**
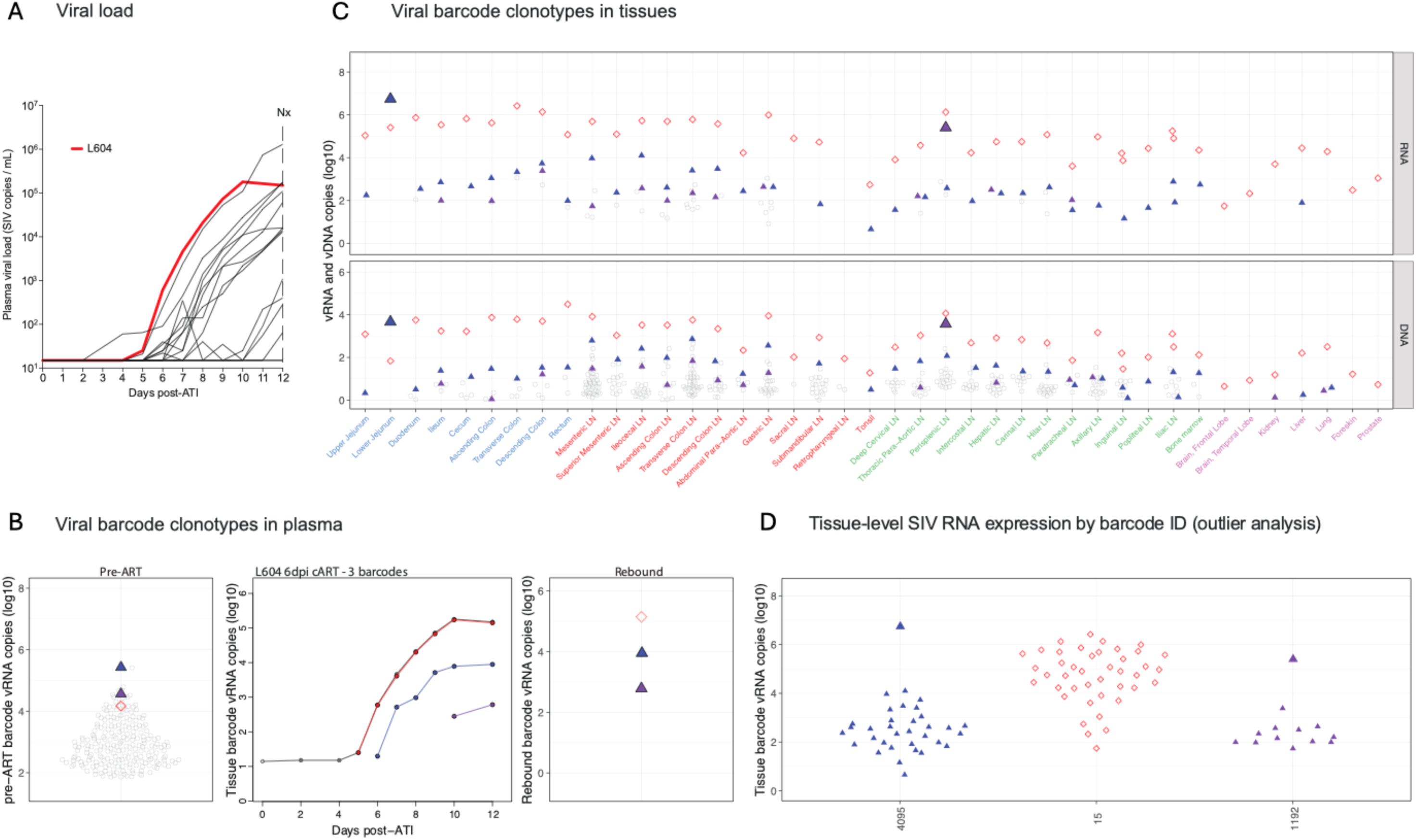
Viral barcode clonotypes in necropsy tissues and contributions to rebound viremia – RM L604, 3 rebounding barcodes. (A) Plasma viral load dynamics following ART discontinuation; the red line indicates the viral load trajectory of RM L604. (B) Left panel: Proportional distribution of viral barcode clonotypes in plasma viremia during primary infection prior to ART initiation, with the 3 barcodes found in rebound plasma highlighted. Middle panel: Calculated rebound viral growth curves of each rebounding barcode lineage with estimated time to a single copy in rebound viremia indicated. Red line indicates the dominant rebounding lineage. Right panel: Proportional distribution of rebound viral barcode clonotypes in plasma at necropsy with barcodes for which a tissue origin site could be identified highlighted. (C) vRNA and vDNA distribution of the three rebounding barcodes (color-coded to panel B) in necropsy tissues. No presumptive tissue origin site could be identified for BC.15 (red open symbols). Colored upward-facing triangles represent rebounding barcodes with tissue origin sites (lower jejunum for BC.4095, perisplenic LN for BC.1192) indicated by the large, filled triangles. Grouped tissue categories are: GI tract (blue), GI tract draining lymph nodes (red), non-GI lymph tissues (green), non-lymphoid tissue (purple), and blood (black). (D) Viral barcode SIV RNA copies in tissue, grouped by barcode ID. Each point represents an individual barcode detected in the indicated tissue.

Animal L681 also rebounded on day 6 following ART discontinuation, with 22 barcodes identified in rebound viremia of 170,000 copies/mL at necropsy on day 12 following ART discontinuation (**Fig. 5A**). All the barcodes in rebound plasma viremia were in the upper half of the primary viremia plasma barcode distribution prior to ART initiation (**Fig. 5B**). Barcode tissue analysis showed widespread distribution of rebounding lineages in the GI tract and GI-associated LNs, and other lymphoid tissues (**Fig. 5C**). The initial barcode detected in rebound plasma (red, BC.4653) was too widely distributed across tissues by day 12 to determine a presumptive tissue origin site. However, outlier analysis of 10 other secondary rebound barcode lineages allowed inference of tissue origin sites, including the duodenum for the third highest rebounding barcode at necropsy (blue, BC.2154) and two additional secondary rebounding lineages (purple, BC.25; pink, BC.2791), as well as a gastric LN for two more secondary rebounding barcodes (teal, BC.3084; green BC.2636). Identification of multiple rebound barcodes originating from specific individual tissue specimens suggests that local tissue factors may be critical in providing the microenvironment necessary for tissue replication leading to viral rebound.^12^

**Figure 5.**
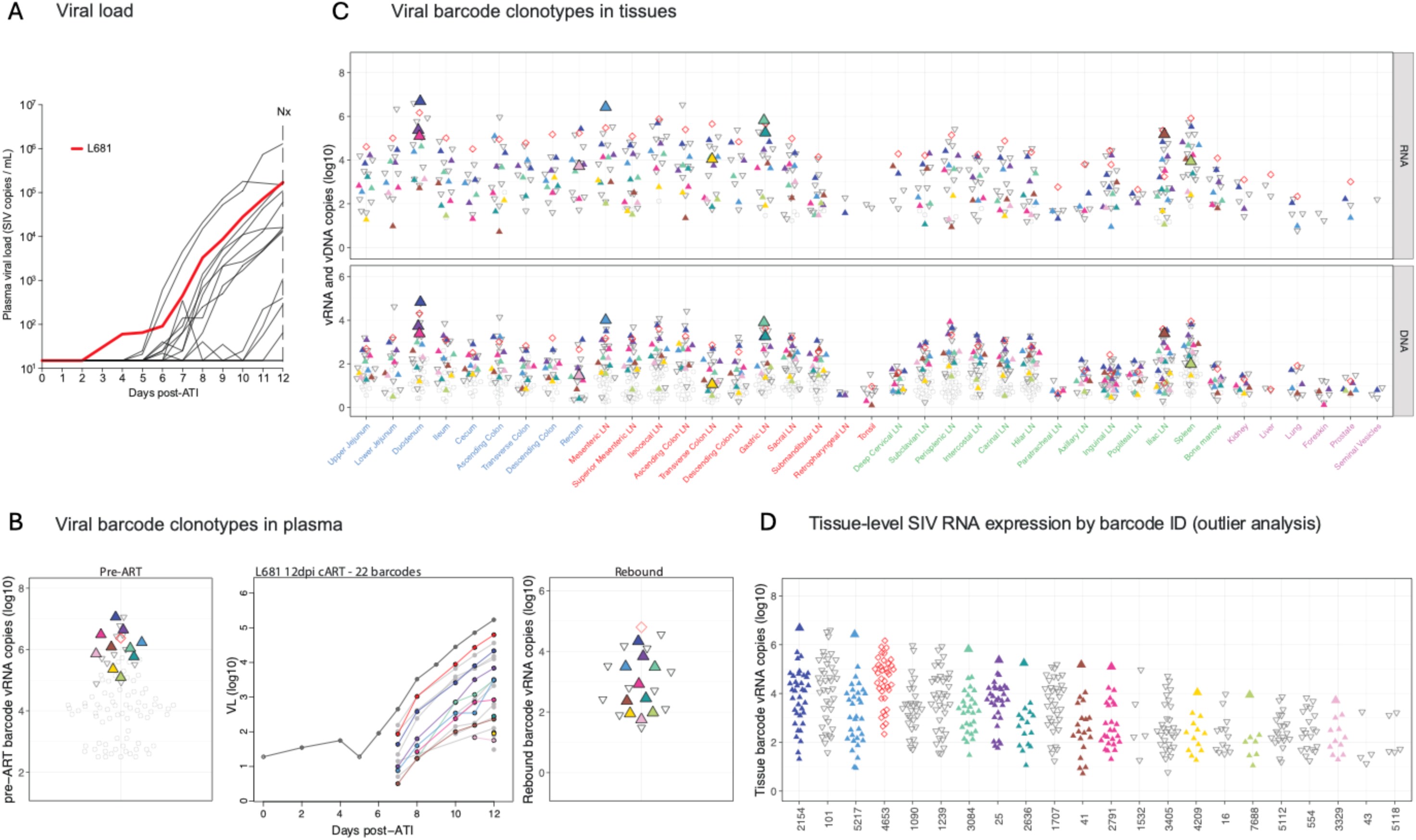
Viral barcode clonotypes in necropsy tissues and contributions to rebound viremia –RM L681, 22 rebounding barcodes. (A) Plasma viral load dynamics following ART discontinuation; red line indicates the viral load trajectory of RM L681. (B) Left panel: Proportional distribution of viral barcode clonotypes in plasma viremia during primary infection prior to ART initiation, with barcodes found in rebound plasma highlighted. Middle panel: Rebound viral growth curves of each rebounding barcode lineage with estimated time to a single copy in rebound viremia indicated. Red line indicates the dominant rebounding lineage, BC.4653. Grey lineages correspond to clones detected in rebound plasma but without an identified presumptive tissue origin site. Right panel: Proportional distribution of rebound viral barcode clonotypes in necropsy plasma with barcodes for which a tissue origin site could be identified highlighted. (C) vRNA and vDNA distribution of all rebounding barcodes (color-coded to panel b) in necropsy tissues. Open symbols indicate barcodes without tissue origin site detected. Colored upward-facing triangles represent rebounding barcodes with a tissue origin site indicated by the large symbol. Grouped tissue categories are: GI tract (blue), GI tract draining lymph nodes (red), non-GI lymph tissues (green), non-lymphoid tissue (purple), and blood (black). (D) Viral barcode SIV RNA copies in tissue, grouped by barcode ID. Each point represents an individual barcode detected in the indicated tissue.

The remaining RMs exhibited 5-38 barcodes in rebound plasma viremia, and outlier analysis inferred sites for tissue origins for a subset of these rebounding barcodes (**Fig. S7-17**). In total, presumptive tissue origin sites were identified for 56 of the 175 total barcode lineages documented in rebound plasma from the 14 viremic RMs following ART discontinuation. Of these, 20 tissue origin sites were found in the GI tract, 24 in GI-associated LNs, and 12 in non-GI lymphoid tissues (**Fig. 6A**). To test if there was a significant enrichment of rebound origin sites based on tissue type, we used mixed-effects logistic regression to assess if tissue group was a significant predictor of viral rebound among the 380 distinct tissue specimens from the 11 viremic RMs with at least one detectable tissue origin site. The odds of contributing barcodes to rebound viremia were 2.6-fold higher for GI tissues (*p*=0.026) and 3.1-fold higher for GI-associated LNs (*p*=0.004) relative to non-GI lymphoid tissues (**Table S3C**). Additionally, GI tissues had the greatest number of tissue sites with more than one inferred tissue origin barcode, including two tissues containing three tissue origin barcodes and a third site with two tissue origin barcodes (**Fig. 6B**). These data illustrate a continuum of viral rebound following ART discontinuation from no rebound to a single barcode lineage rebounding to multiple barcode lineages rebounding, as well as progression from focal tissue viral replication to systemic dissemination with tissue reseeding of replicating lineages. Our analyses also implicate the GI tract and GI-associated LNs as the predominant anatomic sites for the origin of viral rebound following ART discontinuation, suggesting that local tissue microenvironments may impact the probability of a given provirus to contribute to rebound viremia.

**Figure 6.**
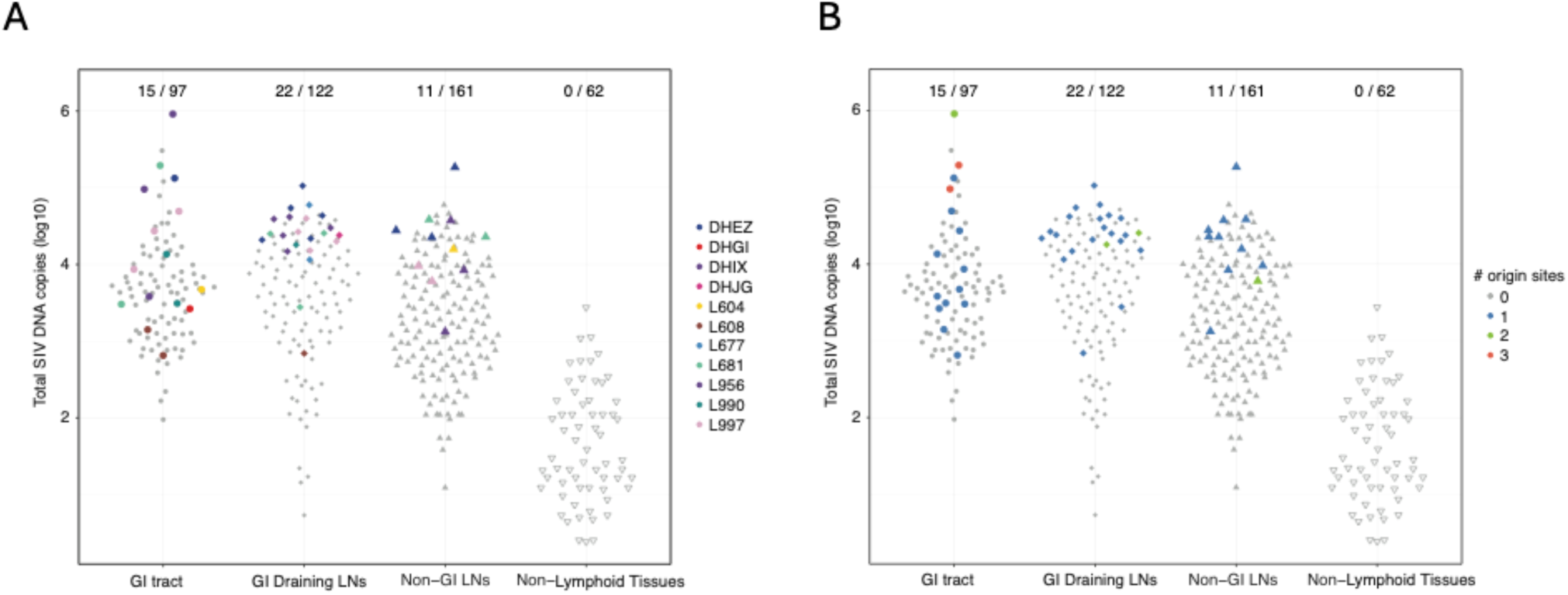
Summary of tissue origin sites contributing to viral rebound. (A) Number of barcode defined rebound origin sites per RM, stratified by anatomic tissue group. Each point represents the total log₁₀ SIV DNA copies per tissue group (GI tract, GI-draining LNs, non-GI LNs, and non-lymphoid tissues). (B) Total number of barcodes originating from individual tissue samples. Fractions indicate the number of tissue origin sites identified out of the total examined.

## Transcriptomic signatures of viral rebound following ART discontinuation

We hypothesized that the early replication events within tissues leading to rebound viremia following ART discontinuation may be associated with systemic changes that may be detectable by transcriptomic signatures in peripheral blood. To characterize potential peripheral blood signatures of viral rebound, we performed bulk whole blood RNA sequencing at the time of ART discontinuation on day 0, day 2, and daily from day 4-12 following ART discontinuation. Transcriptomic analyses were aligned to the time when plasma SIV RNA first exceeded the commonly used clinical threshold of 50 copies/mL with sustained increases thereafter. Gene set enrichment analysis (GSEA) was conducted on differentially expressed genes (DEGs) among all rebounders, comparing transcriptomic profiles prior to ART discontinuation with the sampling timepoints following ART discontinuation but prior to plasma viremia >50 copies/mL (**Fig. S18**). This transcriptomic analysis revealed upregulation of T cell, proinflammatory, cytokine, and innate immune pathways (**Fig. 7A, Fig. S19A**), as well as pathways related to metabolism, cell cycle, and chromatin remodeling (**Fig. 7B, Fig. S19B**; GSEA: false discovery rate [FDR] *q*<0.25) prior to detectable rebound plasma viremia. We assessed the kinetics of transcriptomic changes over time, identifying upregulation of key pathways, including gamma delta T cells, IRF7 activation, TLR signaling, T cell differentiation, metabolism, and cell cycle as early as 6-9 days before rebound plasma viremia of >50 copies/mL was reached (**Fig. 7C-D**). Individual genes activated included canonical T cell signaling components (*LCK*, *CD3D*, *CD247*, *ITM2A*), MHC class I antigen presentation mediators (*CALR*, *PDIA3*), cytotoxic granule proteins (*GZMB*, *GZMK*), and regulators of inflammatory signaling (*JAK1*, *JAK2*, *STAT1*, *MAP2K6*, *FASLG*). These findings suggest the onset of coordinated adaptive and innate immune responses prior to detectable viremia, consistent with peripheral detection of tissue-level viral activity (**Fig. 7E-F**).

**Figure 7.**
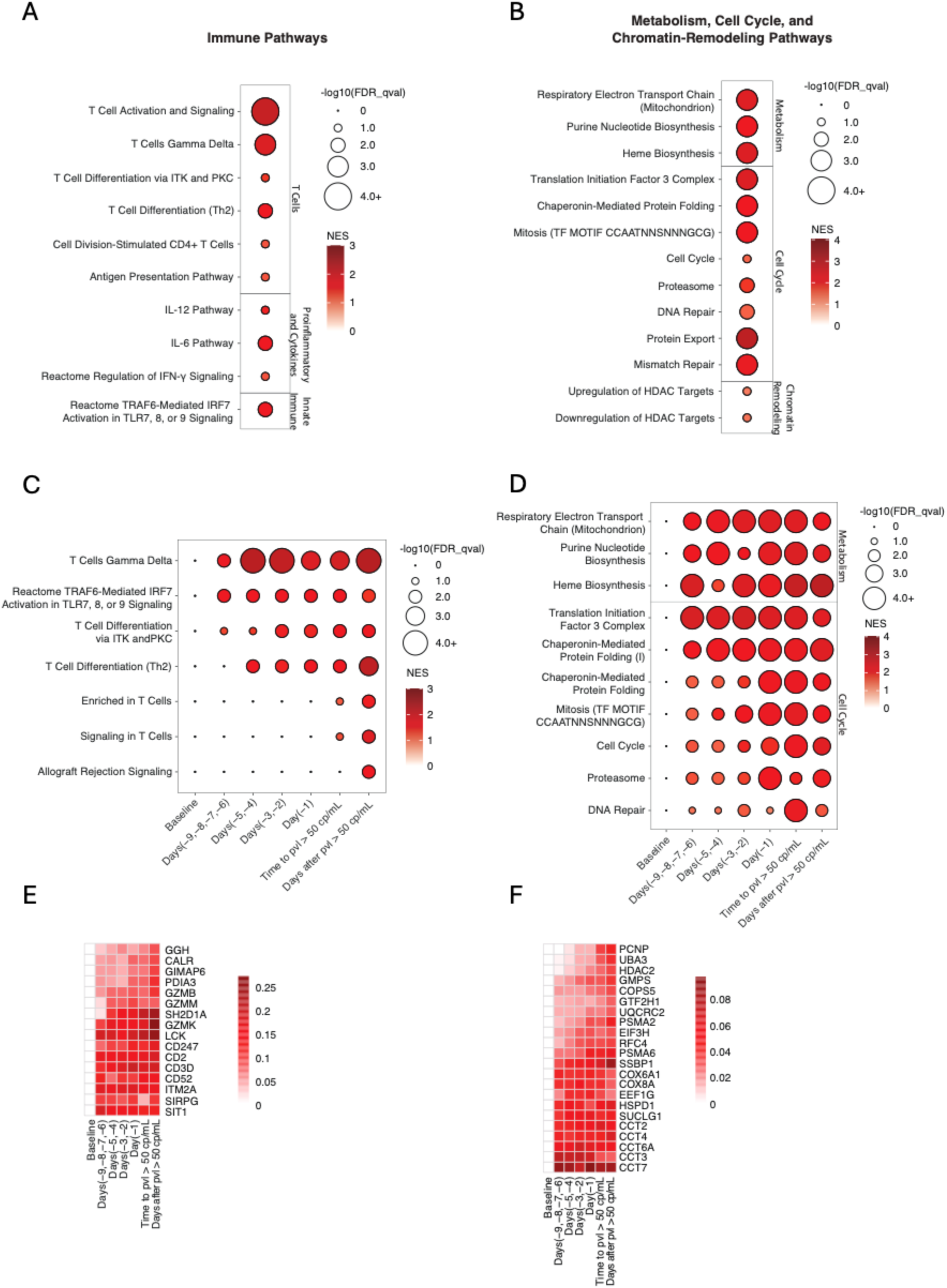
Transcriptomic changes following ART discontinuation. Upregulated transcriptomic pathways compared to baseline (day of ART discontinuation) are shown for samples for the last timepoint following ART discontinuation and prior to rebound viremia <50 copies/mL include (A) immune-related pathways and (B) metabolism, cell cycle, and chromatin remodeling pathways. Time course analysis depicts upregulated pathways relative to baseline for (C) immune pathways and (D) metabolism, cell cycle, and chromatin remodeling pathways across specific timepoints: baseline, days -9 to -6, days -5 to -4, days -3 to -2, and day -1 relative to the first timepoint with rebound viremia >50 copies/mL (defined as day 0 for this analysis). Only significant data (Benjamini-Hochberg-adjusted *p*<0.05) are shown. Circle size reflects significance; color gradient represents the normalized enrichment score (NES). (E) and (F) time course of genes corresponding to consolidated pathways identified in (C) and (D), respectively.

To assess if the upregulation of these pathways represented the early effects of viral replication in tissues following ART discontinuation or alternatively was related to the metabolic or other effects of discontinuation of antiretroviral drugs, we performed a control experiment using a previously published cohort of SIVmac251-infected, ART-suppressed RMs that initiated ART on day 0, day 1, day 2, or day 3 following SIV inoculation.^13^ In this prior study, ART was administered for 24 weeks and was then discontinued. Of the 20 animals, 9 animals exhibited viral rebound following ART discontinuation, and 11 animals did not rebound and were presumed to be uninfected as a result of post-exposure prophylaxis. We performed bulk RNA sequencing in this SIV-uninfected cohort following ART discontinuation and compared GSEA signatures with the SIV-infected cohort. These data suggest that the transcriptomic signatures related to metabolism, cell cycle, chromatin remodeling (**Fig. 8A**), and immune responses (**Fig. 8B**) following ART discontinuation in SIV-infected animals were associated with resumption of viral replication in tissues and impending viral rebound.

**Figure 8.**
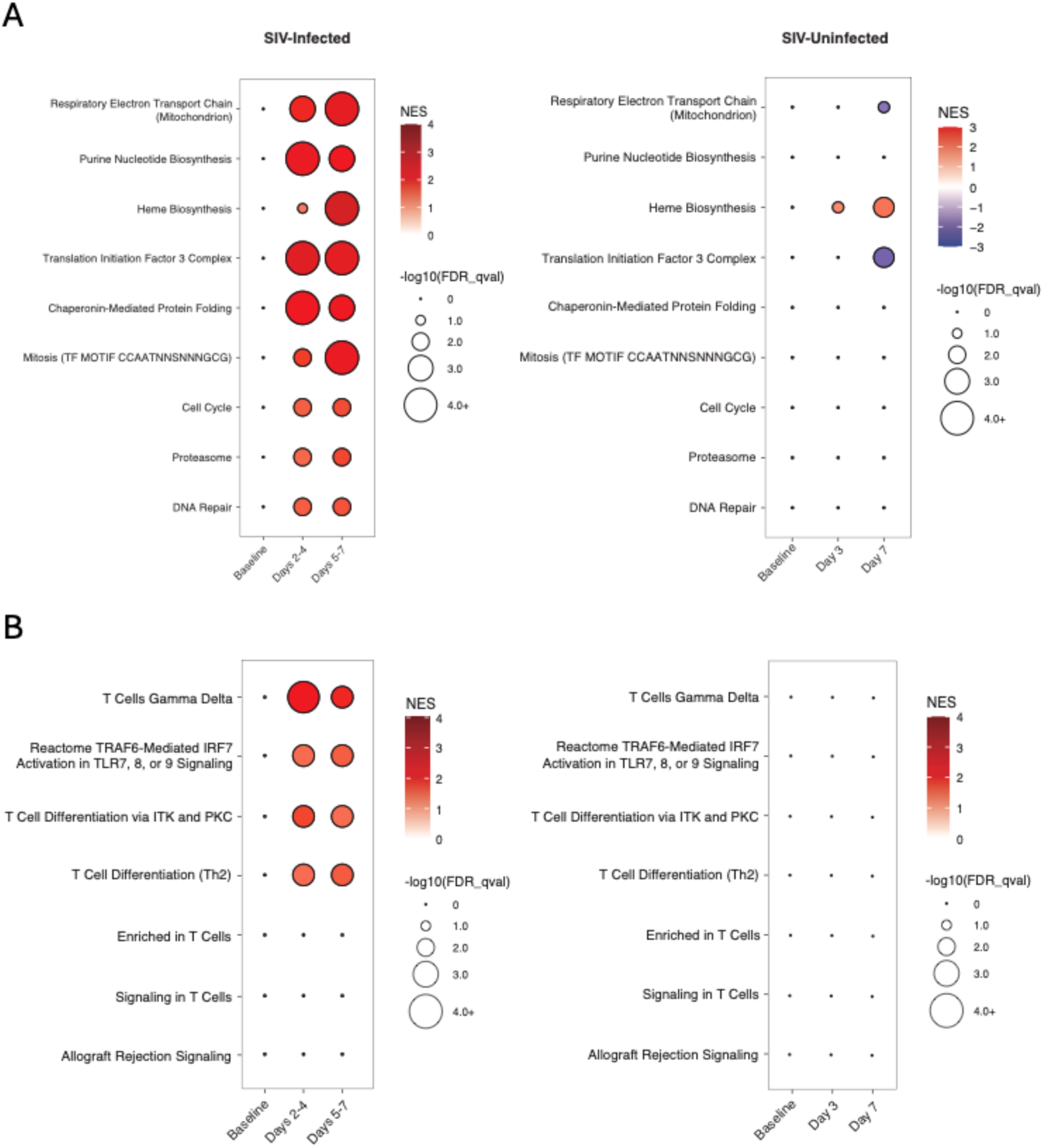
Peripheral blood biomarkers following ART discontinuation in SIV-uninfected RMs. Time course analysis of (A) metabolism, cell cycle, and chromatin remodeling pathways and (B) immune pathways in the above cohort of SIV-infected RMs and a separate cohort of SIV-exposed but uninfected RMs following ART discontinuation. Only significant data (Benjamini-Hochberg-adjusted *p*<0.05) are shown. Circle size reflects significance; color gradient represents the normalized enrichment score (NES), with red indicating upregulation and blue indicating downregulation.

Linear regression analysis of gene expression on the day of ART discontinuation revealed that increased time to rebound was associated with pathways involved in metabolic regulation and cellular biosynthesis, including heme biosynthesis (*r*=0.72, *p*=0.002), naïve B cell surface signature (*r*=0.65, *p*=0.007), drug metabolism (*r*=0.71, *p*=0.002), and cardiomyocyte differentiation via BMP receptors (*r*=0.75, *p*<0.001), while shorter time to rebound correlated with proteasome activity (*r*=-0.71, *p*=0.002) and antigen presentation (*r*=-0.78, *p*<0.001) pathways (**Fig. S20A-B**). Similarly, Cox proportional hazards analysis identified proteasome activity as a predictor of shorter rebound time and metabolic pathways as indicators of increased time to rebound (**Fig. S20C**). These findings suggest that immune activation and cellular degradation mechanisms are active shortly after initial viral replication in tissues and prior to plasma viremia.

## Plasma proteomics

We finally performed plasma proteomic profiling using mass spectrometry at various timepoints relative to baseline. T-tests identified significant upregulated and downregulated proteins over time, and enrichment analysis using the Fisher Exact Overlapping Test was applied to determine the overlap of upregulated proteins within the same pathway sets analyzed in the bulk RNA sequencing (**Figs. 7-8**). Consistent with the transcriptomic data, we observed upregulation of T cell signaling, proinflammatory cytokines, and innate immune pathways (**Fig. S21A**), as well as metabolism, cell cycle, and chromatin remodeling pathways (**Fig. S21B**), prior to plasma viral rebound. Temporal analysis of proteomic pathways following ART discontinuation revealed early upregulation (4-9 days prior to rebound viremia) of proinflammatory and cytokine pathways, including IL-12, IL-6, IL-1, and IL-8 signaling, interferons, and innate immune pathways such as macrophages and dendritic cell activation (**Fig. S21C**) as well as metabolic signatures (**Fig. S21D**).

## Discussion

We characterized the viral dynamics of rebound viremia following ART discontinuation in RMs, using the barcoded virus SIVmac239M to track the processes at the level of individual barcode clonotypes and to identify presumptive tissue origin sites for a subset of the barcodes identified in rebound plasma viremia. Animals demonstrated a broad spectrum of rebound outcomes at necropsy from no detectable viral rebound to full rebound with plasma viral loads >10^6^ SIV RNA copies/mL. We used extensive tissue sampling, vRNA and vDNA quantification, and barcode analyses along with a modified outlier analysis to identify tissue sites with elevated levels of individual barcode vRNA at necropsy as presumptive tissue origin sites for 56 of 175 barcode lineages contributing to rebound viremia. Moreover, we demonstrated inflammatory and metabolic changes in peripheral blood prior to detectable rebound viremia, presumably reflecting viral replication in tissues.

Our data suggest a model in which individual viral clonotypes that contribute to viral rebound originate from serial oligofocal events, primarily in GI tissues and GI-draining LNs, with initial local tissue spread followed by systemic spread. These findings have important implications for our understanding of the pathogenesis of viral rebound and for the development of next generation HIV-1 cure strategies. Our results, focused on later stages of viral rebound with necropsies performed on day 12 following ART discontinuation and employing an alternative analytical approach suited to this context, reinforce those of a companion study using the same RM/SIVmac239M model^12^ which focused on earlier necropsy timepoints (day 5 and day 7 following ART discontinuation) to define the earliest events in tissues leading to viremic rebound and to identify initial rebound barcodes and their anatomic origins, 96% of which were found in GI tract and GI associated LNs. These two studies thus underscore the critical role of the GI tract and associated lymphoid tissue microenvironments in viral rebound, including both the initial clonotype lineages and the subsequent clonotype lineages that contribute to rebound, even in the context of systemic inflammation and immune activation.^14^

Barcodes with higher primary infection levels prior to ART initiation generally correlated with those found more frequently in rebound plasma viremia after ART discontinuation. These findings suggest that early viral replication dynamics for individual viral barcode clonotypes can impact viral seeding of tissues during acute infection and eventually contribute to viral rebound. However, while more highly represented barcodes had a higher probability of contributing to viral rebound, the most highly represented barcode in both primary infection and in tissues did not always initiate rebound viremia, suggesting the importance of additional contributing factors, such as the cellular and local tissue microenvironment.^1,7,15^

Multi-omic profiling of peripheral blood showed early upregulation of immune activation and metabolic pathways prior to detection of rebound viremia and likely reflecting viral replication in tissues. These findings align with previous studies showing early upregulation of interferon-stimulated genes, antiviral restriction factors, and inflammatory responses following ART discontinuation.^16^ Metabolic shifts, including glycolysis and mitochondrial function, suggest a systemic response to viral replication in tissues before virus in blood can readily be measured. Notably, the proteomic signatures identified in our RM model closely parallel data from recent human studies of ART discontinuation, in which individuals with impending viral rebound exhibited increased immune activation, interferon signaling, and apoptosis, while virologic controllers displayed metabolic shifts and reduced inflammation.^17,18^ Given that these molecular signatures emerged prior to detectable plasma viremia, they likely reflected systemic inflammation and metabolic reprogramming triggered by viral expression and local viral replication in tissues. These signatures could inform the development of biomarkers to predict or monitor viral rebound and to guide therapeutic approaches to target these processes for HIV-1 cure strategies.

Our study has several limitations. First, we were unable to identify the origin sites of all clonal lineages due to substantial viral replication in most animals by day 12 following ART discontinuation. Second, although we evaluated multiple tissues at necropsy, these tissues still only reflected a fraction of the total GI and lymphoid tissues in the animals. Third, our transcriptomic and proteomic profiling was limited to peripheral blood and may not reflect the microenvironment in relevant tissues. Fourth, although the barcoded SIV model allows high-resolution clonal lineage tracking, extrapolation to HIV-1 in humans requires cautious interpretation.

Our findings indicate that viremic rebound following ART discontinuation is initially driven by oligoclonal and oligofocal replication of individual viral clonotypes, predominantly in GI tissues and GI-associated LNs, leading to systemic dissemination with reseeding of tissues with these initial rebounding clonotypes, as well as serial initiation of viral replication in tissues by additional viral clonotypes. The early events following ART discontinuation in tissues drive upregulation of inflammatory and metabolic pathways that can be detected in peripheral blood prior to plasma viremia, offering mechanistic insights and potential biomarkers of impending plasma viral rebound. Future studies should define the molecular and immunologic factors within permissive tissue niches that enable initial viral reactivation, which would inform HIV-1 cure strategies that aim to target these processes.

## Author Contributions

D.H.B., L.J.P., J.D.L., and B.F.K designed the study. I.V.K., J.D.L., B.F.K., V.E.K.W.-S., and D.M.T. performed and analyzed virologic and immunologic assays. B.F.K., E.K., T.T.I., C.A.G., A.C and C.M.F. performed and analyzed barcode sequencing. I.V.K., M.A., and A.C. performed the transcriptomic and proteomic analyses. R.G. provided the ART regimen. W.J.R. and M.J.F. led the clinical care of the macaques. I.V.K., J.D.L., B.F.K., D.H.B., and all co-authors contributed to the writing and editing of the paper.

## Acknowledgments

We thank BIDMC’s Genomics and Proteomics Core, the NHP Genomics Core at Emory Yerkes National Primate Research Center, the Pacific Northwest National Laboratory (PNNL), the Quantitative Molecular Diagnostics Core, Viral Evolution Core, and Computational Virology Group of the AIDS and Cancer Virus Program of the Frederick National Laboratory for Cancer Research, Tetyana Murzda, and Tiffany Markel. This project was supported by the Gates Foundation (INV-002377 to D.H.B., L.J.P., J.D.L.) and the National Institutes of Health (UM1AI164556, P01AI177687, P01AI169615, R01AI149670 to D.H.B.; UM1AI164560 to L.J.P.; and 75N91019D00024 to B.F.K., J.D.L.) The content of this publication does not necessarily reflect the views or policies of the Department of Health and Human Services, nor does mention of trade names, commercial products, or organizations imply endorsement by the U.S. Government. The funders contributed to the study design but were not involved in the study operations, data collection, data analysis, data interpretations, decision to publish, or preparation of the paper. The data and statistical leads had access to all the data. D.H.B. had final responsibility for the decision to submit for publication.

## STAR★Methods

### RESOURCE AVAILABILITY

#### Lead Contact

Further information and requests for resources and reagents should be directed to and will be fulfilled by the Lead Contact, Dan H. Barouch.

#### Materials Availability

This study did not generate new unique reagents. The molecularly barcoded SIVmac239M construct and the intact proviral DNA assay (IDPA) primers and probes are available upon request, subject to institutional approvals and standard material transfer agreements.

#### Data and Code Availability

- Bulk RNA sequencing data generated in this study are available at the NCBI Gene Expression Omnibus (GEO) under accession number **GSE294867**.
- Viral load measurements and barcode analyses are included in Data Source files accompanying this manuscript.
- Additional requests for data supporting the findings of this study are available from the Lead Contact upon reasonable request.
- Custom code used for clustering and outlier detection is available from the corresponding author upon request.

### EXPERIMENTAL MODEL AND STUDY PARTICIPANT DETAILS

#### Animals

Eighteen outbred, Indian-origin adult male and female RM (*Macaca mulatta*), housed at AlphaGenesis (Yemassee, SC), were infected intravenously at week 0 with 5,000 IU SIVmac239M, as previously described.^6^ Animals were randomly assigned to one of three groups and were started on ART at either day 6, 9, or 12 following challenge to allow different levels of RCRV seeding. Biopsy sampling prioritized lymph nodes and GI tissues. At week 70, ART was discontinued and RMs were necropsied on day 12 following ART discontinuation. Blood was collected on days 0, 2, and 4-12 after ART discontinuation, and at necropsy tissues were collected from all major organ systems on day 12. Immunologic and virologic assays were performed blinded. Two RMs were euthanized for procedure-related complications and were not followed through the ART discontinuation phase of the study. All RMs were housed and cared for under protocols approved by the Institutional Animal Care and Use Committee (IACUC). Prior to enrollment, animals were screened and confirmed to be free of simian retrovirus D, STLV-1, Herpes B, and Mycobacterium tuberculosis. Major histocompatibility complex (MHC) class I genotyping was performed to exclude the presence of protective alleles (Mamu-A*01, -B*08, -B*17). Demographics are provided in Table S2.

#### Ethics statement

All the work described in this manuscript adhered to the guidelines outlines in National Institutes of Health (NIH) Guide to the Care and Use of Laboratory Animals. All studies were approved by the AlphaGenesis Institutional Animal Care and Use Committee (IACUC) and was conducted in accordance with federal, state, and local laws and regulations.

### METHOD DETAILS

#### Antiretroviral Therapy (ART) regimen

The pre-formulated antiretroviral therapy (ART) cocktail was provided by Gilead Sciences (Foster City, CA) and contained 5.1 mg/mL of tenofovir disoproxil fumarate (TDF), 40 mg/mL of emtricitabine (FTC), and 2.5 mg/mL of dolutegravir (DTG) dissolved in 15% (v/v) kleptose,^2^ which was adjusted to a pH of 4.2. The ART cocktail was administered daily to the study animals by subcutaneous injection at 1 mL/kg body weight for a period of 70 weeks.

#### Plasma Viral Load Assays

Plasma SIV vRNA was measured essentially as described^19^ at multiple time points throughout duration of the study, including prior to the initiation of ART and after the discontinuation of ART using a gag targeted real time assay with a threshold of 15 SIV RNA copies/mL.^20^

#### Quantitative Evaluation of Cell-Associated vDNA and vRNA

For PBMC and tissue specimens, levels of vRNA and vDNA were measured essentially as described using assays targeted to gag.^21,22^

#### Intact Proviral DNA Assay (IPDA)

To isolate CD4+ cells from frozen PBMCs, we utilized the EasySep NHP CD4+ T Cell Isolation Kit (Stem Cell), and total cellular DNA was extracted from the isolated CD4+ cells using the QIAmp DNA Blood Mini Kit (Qiagen). DNA concentration was measured by NanoDrop 2000 (ThermoFisher Scientific). To quantify intact proviral SIV DNA, we performed the SIV-specific digital droplet PCR (ddPCR) intact proviral DNA assay as described previously.^23^ Briefly, up to 300 ng sample DNA was added to a master mix containing 2x ddPCR Supermix for Probes (no dUTP, Bio-rad), 600 nM of each primer, and 200 nM of each probe for each 22 uL reaction. Cell equivalents were determined by quantifying copies of macaque RPP30 in parallel, allowing for the absolute quantification of intact proviral SIV DNA. ddPCR outputs were analyzed using QuantaSoft Analysis-Pro (Bio-Rad).^24^

#### Barcode Sequencing and Enumeration

To characterize barcode clonotype populations in plasma during primary SIV infection prior to ART initiation, during off-ART rebound viremia, and from vRNA positive PBMC and biopsy and necropsy tissue specimens, barcode sequence analysis was performed on viral nucleic acid samples as previously described^10^ including both vDNA and vRNA (cDNA) sequencing for PBMC and tissue samples.

#### Outlier Analysis to Identify Tissue Origin Sites of Rebounding Barcode Lineages

To distinguish the rebound origin site for an individual rebounding lineage expressing vRNA in multiple different tissues, we used Density-Based Clustering of Applications with Noise (DBSCAN) to cluster tissues based on the barcode vRNA and vDNA levels to identify a single outlier site with elevated vRNA expression that did not cluster with any other tissues. The analysis was performed using the “DBSCAN package” in R (version 4.1.1) for each rebounding barcode with vRNA and vDNA in more than 6 samples. The algorithm iteratively clusters points into closely packed, dense regions, separated by regions of lower density, and identifies outliers without assigning them to any cluster. The minimum number of points required to form a dense cluster was chosen as *k*=3 for one-dimensional analysis (based on vRNA only) and *k*=4 for two-dimensional analysis (based on vRNA vs. vDNA). For each barcode, the k-nearest neighbors (kNN) distance method was used to determine *epsilon,* the maximum distance between two points for them to be assigned to the same cluster. The kNN-distance (total distance of a point to its k-nearest neighbors) was calculated for each point, which were then rank-ordered, resulting in a generally piecewise linear relationship with a single-breakpoint (defining *epsilon*), which we estimated automatically by fitting a piecewise linear regression using the “Segmented” package in R. Using this process, we identified all samples that were not assigned to a cluster (vRNA or vRNA vs. vDNA) for each rebounding barcode lineage (with vRNA+ and vDNA+ in at least 7 samples), designated as cluster-outliers. We then compared vRNA expression in these cluster-outliers to non-outlier tissues to identify sites with elevated local replication compared to background vRNA levels or replication in secondary sites. For each lineage, we rank-ordered all samples based on their vRNA levels and defined the sample with the highest vRNA level as the putative rebound origin if (1) it was a cluster-outlier and (2) its vRNA (log_10_) distance to the next-highest sample exceeded a threshold level η associated with disseminated replication or background vRNA expression, defined as the 99^th^ quantile (η=0.52 log_10_) of the distribution of vRNA differences between non-outlier samples across all lineages. The origin sites were categorized into anatomical groups to assess patterns of tissue-specific initiation of viral rebound.

#### Regression Analyses

All regression analyses were performed in R using the lme4 package for mixed effects (Table S3).

##### Model 1: The effect of pre-ART replication on probability of rebound at the barcode-level

We performed mixed effects logistic regression analyses to investigate for all 16 evaluable animals whether a barcode’s pre-ART peak plasma viral load across was predictive of whether it rebounded (i.e. detected in rebound plasma). Repeated observations within individual animals were accounted for by including a random effect on the intercept (assumed normally distributed with a zero mean).

##### Model 2: The effect of rebound plasma viral loads on barcode vDNA and vRNA levels in tissues

For the 175 rebounding lineages from the 14 viremic animals, we performed mixed-effects linear regression to investigate if the barcode viral load (log_10_) in rebound plasma was predicted by either its total vDNA or vRNA levels (log_10_) across all tissues. Repeated observations within individual animals were accounted for by including a random effect on the intercept (assumed normally distributed with a zero mean).

##### Model 3: The effect of tissue group on probability of rebound at the tissue-level

We performed mixed effects logistic regression analysis to investigate for the 11 animals with at least one identified tissue origin site whether tissue type was predictive of the probability of rebound at the tissue-level. Clustering of observations within individual animals was accounted for by including a random effect on the intercept (assumed normally distributed with a zero mean). The null model (without tissue type as a covariate) was compared to the full model using the likelihood ratio test via ANOVA.

#### Antibody ELISA

SIV-specific antibodies were assessed using a serial Enzyme-Linked Immunosorbent Assay (ELISA) to measure the reactivity of serum antibodies to the SIVmac239 gp160 recombinant protein. To perform the ELISA, serum samples were diluted and added to 96-well plates that were coated with the SIVmac239 gp160 recombinant protein before being washed with Phosphate Buffered Saline (PBS) + 0.05% Tween 20 and blocked with Blocker Casein (Pierce). The plates were incubated, allowing SIV-specific antibodies in the serum to bind to the protein. After another wash to remove unbound antibodies and incubation with rabbit anti-mouse IgG horseradish peroxidase (Thermo Scientific), plates were washed and developed with SureBlue (KPL Laboratories) and stopped with TMB Stop Solution (KPL Laboratories. Plates were read on a VersaMax microplate reader (Molecular Devices) and absorbance at a wavelength of 450nm was recorded. Endpoint titers were determined as the reciprocal of the highest serum dilution that delivered an optical absorbance value above the value of the negative control sera.

#### Cellular Immune Responses

SIV-specific cellular immune responses were assessed using IFN-γ ELISPOT assays. These assays were performed essentially as described, with some modifications to allow for the assessment of cellular immune breadth. To estimate the breadth of virus specific cellular immune responses, PBMC IFN-γ ELISPOT assays were performed using sub-pools of peptides spanning the SIVmac239 Env, Gag, and Pol proteins, providing a simultaneous measurement of T cell responses to multiple epitopes. The limit of detection of this assay was 5 spots/million cells.^9^ Flow cytometric staining was performed to further characterize the phenotype and functional properties of the SIV-specific T cells. This was accomplished utilizing predetermined titers of monoclonal antibodies at concentrations, suggested by the manufacturer (Becton Dickinson), against a panel of surface and intracellular markers, including CD3 (SP34; Alexa Fluor 700), CD4 (OKT4; BV510, Biolegend), CD8 (SK1; APC-Cy7), CD14 (M5E2; BUV737), CD16 (3G8; BV650), CD25 (PE-Cy7; M-A251), CD28 (L293; PerCP-Cy5.5), CD38 (APC; HB-7), CD56 (NCAM16; BV786), CD69 (TP1.55.3; PE-TexasRed; Beckman Coulter), CD95 (DX2; BV711), CCR5 (3A9; PE), CCR7 (3D12; BV421), HLA-DR (BUV-395; G46-6), Ki67 (B56; FITC), and PD-1 (EH21.1; BV605).

#### Transcriptomic Profiling

Transcriptomic Bulk RNA sequencing analysis was conducted on whole blood samples collected in PAXgene tubes, with sequencing by the NHP Genomics Core at Emory Yerkes National Primate Research Center in Atlanta, GA. RNA sequencing was carried out utilizing the Illumina NextSeq 500/550 High Output v2 kits (150 cycles), following the manufacturer’s protocol. The sequencing reads were aligned to the reference genome using the STAR aligner, and downstream differential expression analysis was conducted with the DESeq2 package in R. This analysis identified normalized expression counts and differentially expressed genes (DEGs) between timepoints post-ATI compared to baseline (the day of treatment interruption), with statistical significance defined by adjusted *p*-values (Benjamini-Hochberg correction) <0.05. Visualization of normalized expression counts was achieved Principal Component Analysis (PCA) and heatmaps. Gene Set Enrichment Analysis (GSEA) was applied to evaluate biological pathways associated with DEGs, leveraging curated pathway sets such as Biological Themes (BTM), C2 pathways, and in-house compiled pathways. Single-cell Linear Expression Analysis (SLEA) was employed to perform a Spearman correlation between gene expression and time to viral rebound, offering insights into how specific genes or pathways may influence the timing of rebound post-ATI. Further, a univariate Cox model was applied to identify pathways significantly associated with time to rebound.

#### Proteomic Profiling

Proteomic analysis was conducted on plasma samples, with sequencing performed at the Pacific Northwest National Laboratory (PNNL) using advanced mass spectrometry techniques to measure a panel of less than 2,000 proteins. Data visualization included the use of Principal Component Analysis (PCA) and heatmaps to depict expression patterns across time points. For statistical analysis, T-tests were employed to compare the levels of upregulated and downregulated proteins at various time points against baseline measurements, identifying significant shifts in protein expression. Additionally, enrichment analysis was conducted using the Fisher Exact Overlapping Test, using the same pathway sets applied in the Bulk RNA sequencing analysis.

### QUANTIFICATION AND STATISTICAL ANALYSIS

Statistical analyses were performed using GraphPad Prism Version 10 (GraphPad Software) and R.

- **Tests used:** Kruskal-Wallis tests for group comparisons; two-sided Spearman correlations for associations; mixed-effects regression for barcode rebound predictors; Cox proportional hazards models for time-to-rebound analyses; Wald tests for barcode distribution comparisons; Wilcoxon rank-sum for tissue vRNA/vDNA comparisons.
- **Sample size (*n*):** For virologic and immunologic data, *n*=number of animals. For barcode-level analyses, *n*=number of unique barcodes detected per animal. For tissue analyses, *n*=number of independent tissue samples.
- **Replicates:** All assays were performed in at least technical duplicate; PBMC and tissue analyses included biological replicates across animals.
- **Center and dispersion:** Data are reported as mean ± SD unless otherwise indicated.
- **Significance thresholds:** Adjusted *p*-values (Benjamini-Hochberg), <0.05 for transcriptomic and proteomic analyses; q<0.25 for GSEA; other tests considered significant at *p*<0.05.
- **Randomization/blinding:** Animals were randomized into ART initiation groups. Assays were performed blinded to treatment group.
- **Inclusion/exclusion:** Two animals were excluded due to unrelated euthanasia during ART. No other exclusions applied.

### ADDITIONAL RESOURCES

No additional resources were generated in this study.

## Supplementary Figure Legends

**Figure S1.**
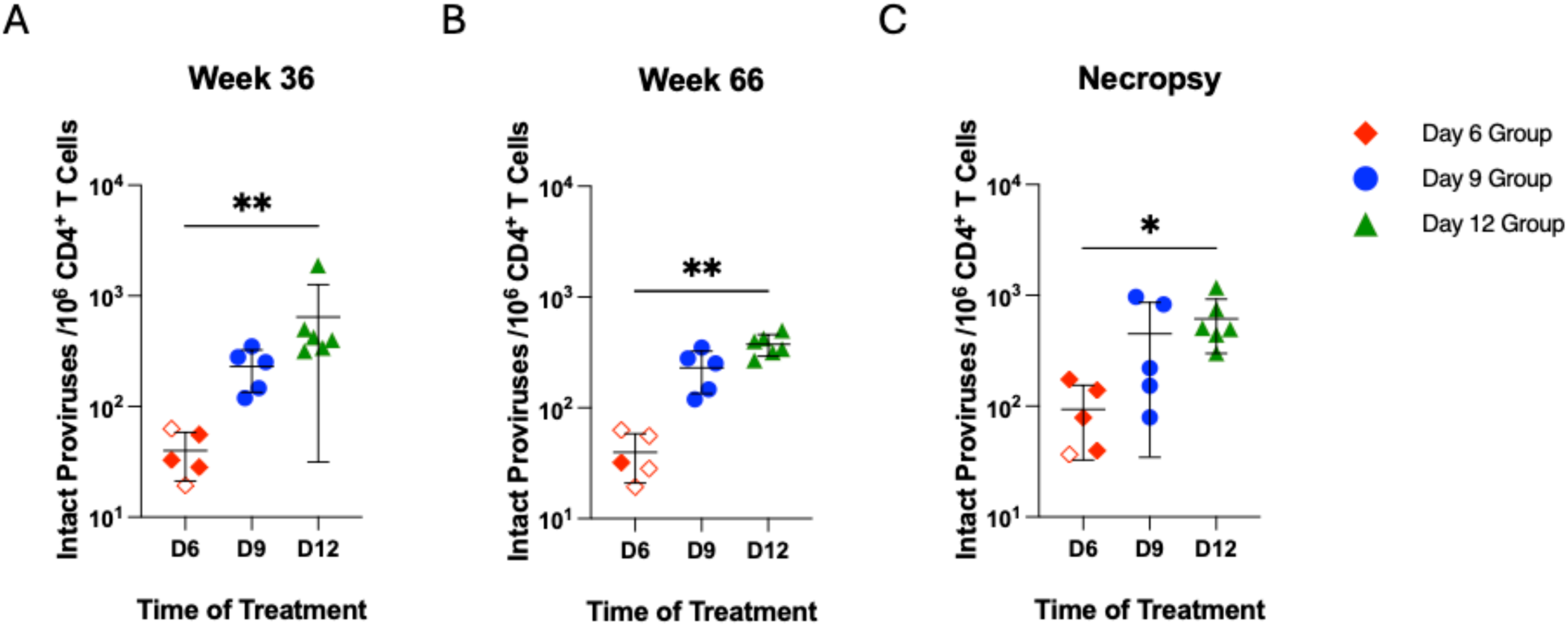
Distribution and quantification of intact proviruses in PBMCs during ART suppression and at necropsy. Quantification of intact proviruses in PBMC using the SIV-specific intact proviral DNA assay (IPDA) at (A) week 36 post-infection, (B) week 66 post-infection, and (C) necropsy. Data is shown for each ART initiation group: day 6 (red diamonds), day 9 (blue circles), and day 12 (green triangles). Black lines indicate mean ± standard deviation. * *p*<0.0332, ** *p*<0.0021. Open symbols are below limit of detection.

**Figure S2.**
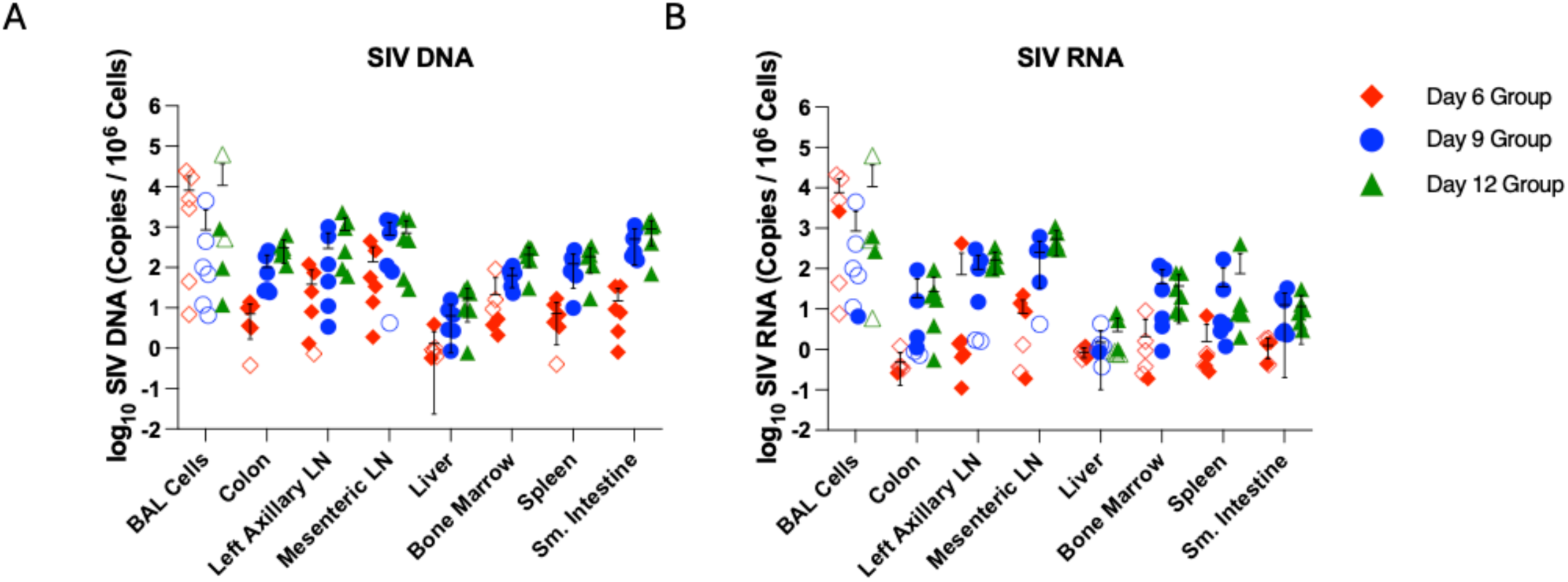
Tissue distribution of cell associated vDNA and vRNA at week 21. Detection of (A) SIV cell-associated DNA (vDNA) and (B) SIV cell-associated RNA (vRNA) in various tissues at week 21 following infection, including in bronchoalveolar lavage (BAL) cells, colon, left axillary LN, mesenteric LN, liver, bone marrow, spleen, and small intestine biopsies. Data is shown for each ART initiation group: day 6 (red diamonds), day 9 (blue circles), and day 12 (green triangles). Black lines indicate mean ± standard deviation. Unfilled circles represent samples in which no replicates were positive for vDNA or vRNA, and an imputed threshold value was plotted to indicate levels below the detection limit.

**Figure S3.**
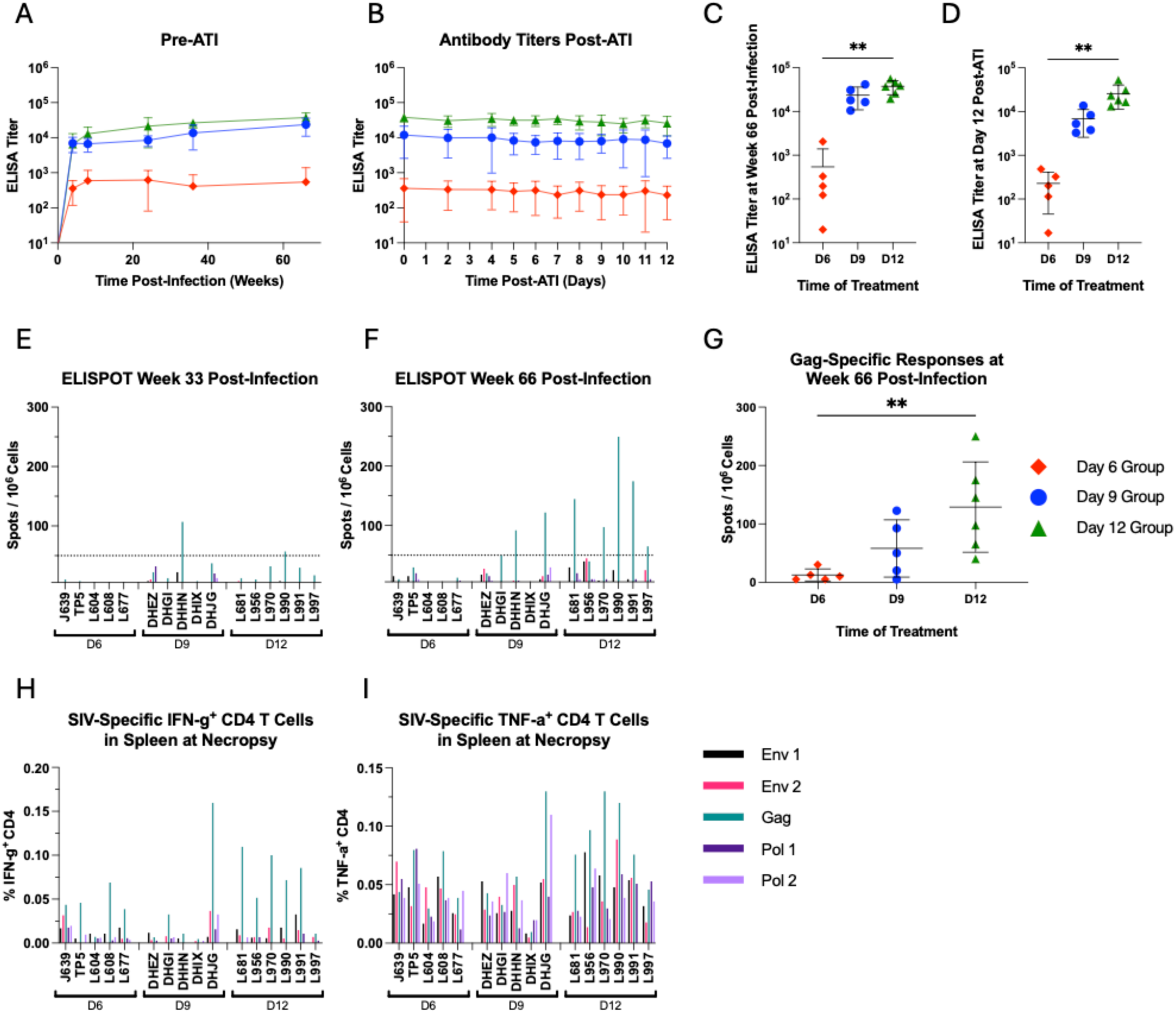
SIV-specific humoral and cellular immune responses during ART and after ART discontinuation. The median titers of serum antibodies to the SIVmac239 gp160 recombinant protein are shown for all treatment groups, day 6 (red), day 9 (blue), and day 12 (green) (A) during ART suppression and (B) after treatment interruption. ELISA antibody titers are shown at (C) week 66 post-infection and (D) day 12 (at necropsy) after treatment interruption. PBMC IFN-γ ELISPOT assays using pools of peptides spanning the SIVmac239 Env, Gag, and Pol proteins at (E) week 33 post-infection and (F) week 66 post-infection, with the dotted line indicating the limit of detection. (G) PBMC Gag-specific cellular immune responses at week 66 following infection. (H) SIV-specific IFN-γ + CD4 T cells in spleen at necropsy and (I) SIV-specific TNF-α + CD4 T cells in spleen at necropsy. Black lines indicate mean ± standard deviation. * *p*<0.0332, ** *p*<0.0021.

**Figure S4.**
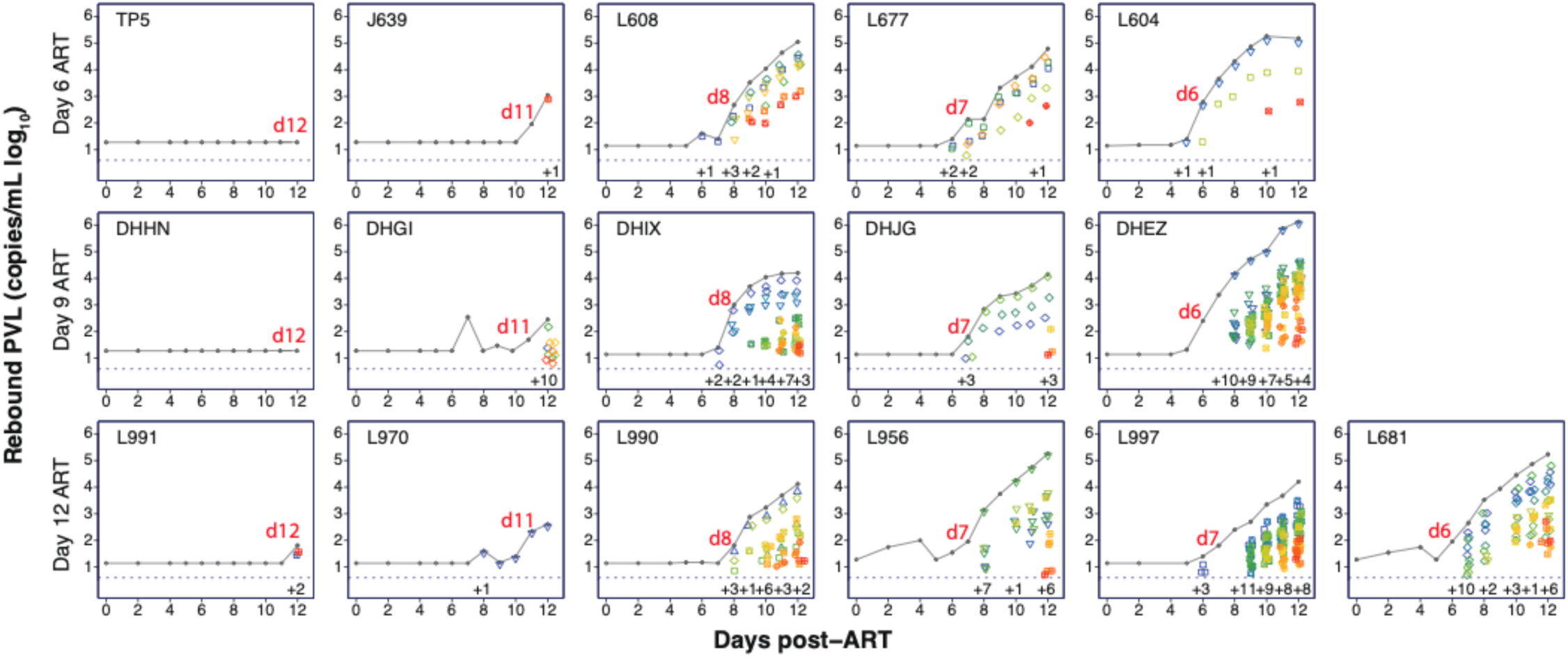
Dynamics of rebound after ART discontinuation. Rebound plasma viral load (RNA copies/mL) from time of ART discontinuation to necropsy for each animal, grouped by ART initiation time. The emergence and proportional contribution to total viral load of distinct barcodes is shown longitudinally, with heatmap color coding of barcodes corresponding to when they were first detected in rebound plasma. For all viremic animals, the number of newly identified barcode lineages per day is shown. The time to plasma viral load rebound of >50 copies/mL is labeled in red as the number of days following ART discontinuation.

**Figure S5.**
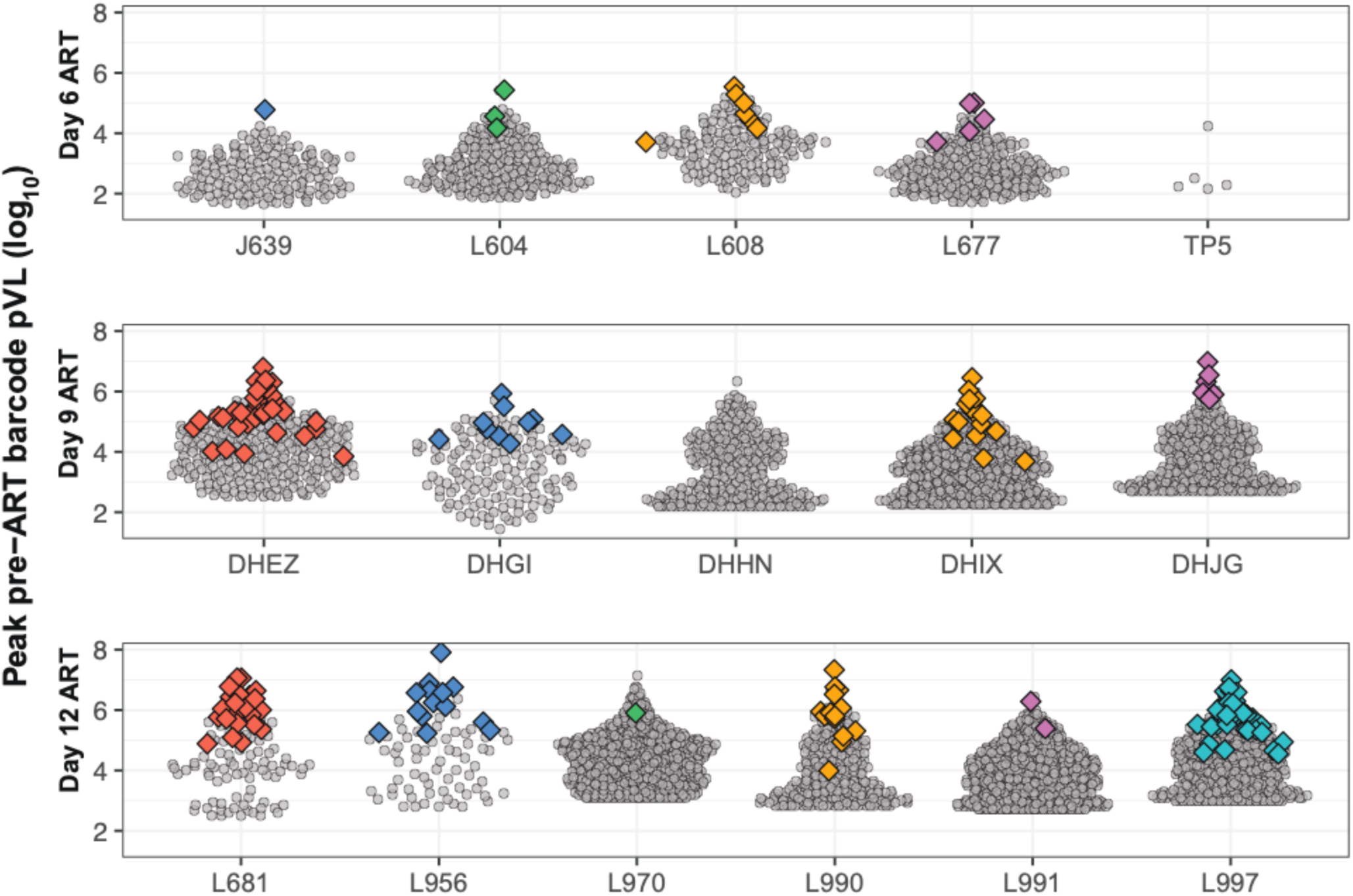
Rebounding barcodes labeled in barcode distribution of plasma viremia during primary infection prior to ART initiation. The barcode plasma viral loads (log_10_ RNA copies/mL) prior to ART initiation is shown for each animal, grouped by ART initiation time. Grey dots correspond to barcodes that were not detected in rebound plasma, whereas colored diamonds indicate rebounding lineages (*n*=175).

**Figure S6.**
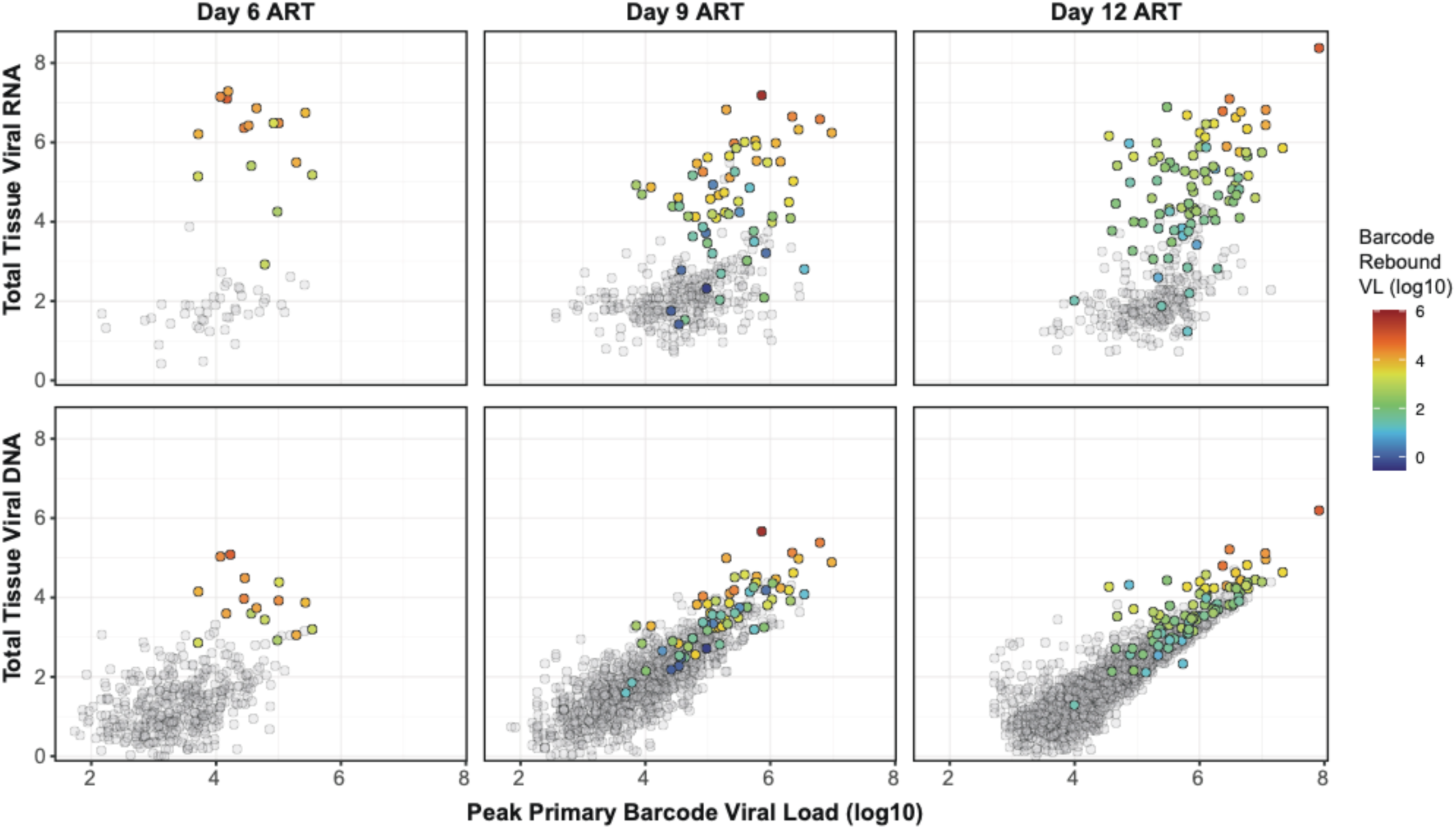
Total barcode vRNA and vDNA levels in necropsy tissues vs. pre-ART and rebound plasma viremia. For animals starting ART at day 6 (left), 9 (middle) and 12 (right), the primary infection viremia prior to ART initation (log_10,_ x-axis) for each individual identified barcode is compared against total vRNA (top) and vDNA (bottom) (log_10,_ y-axis) for that barcode across all analyzed necropsy tissues. Colored dots indicate barcode lineages identified in rebound viremia, with the color scale indicating the level of each individual barcode in rebound viremia (log_10_) at necropsy. Grey dots indicate barcodes that were not detected in rebound plasma viremia. Overall, higher levels of individual rebounding vRNA barcodes in tissues (color) were associated with higher levels in peak primary infection viremia and higher representation in rebound viremia.

**Figure S7.**
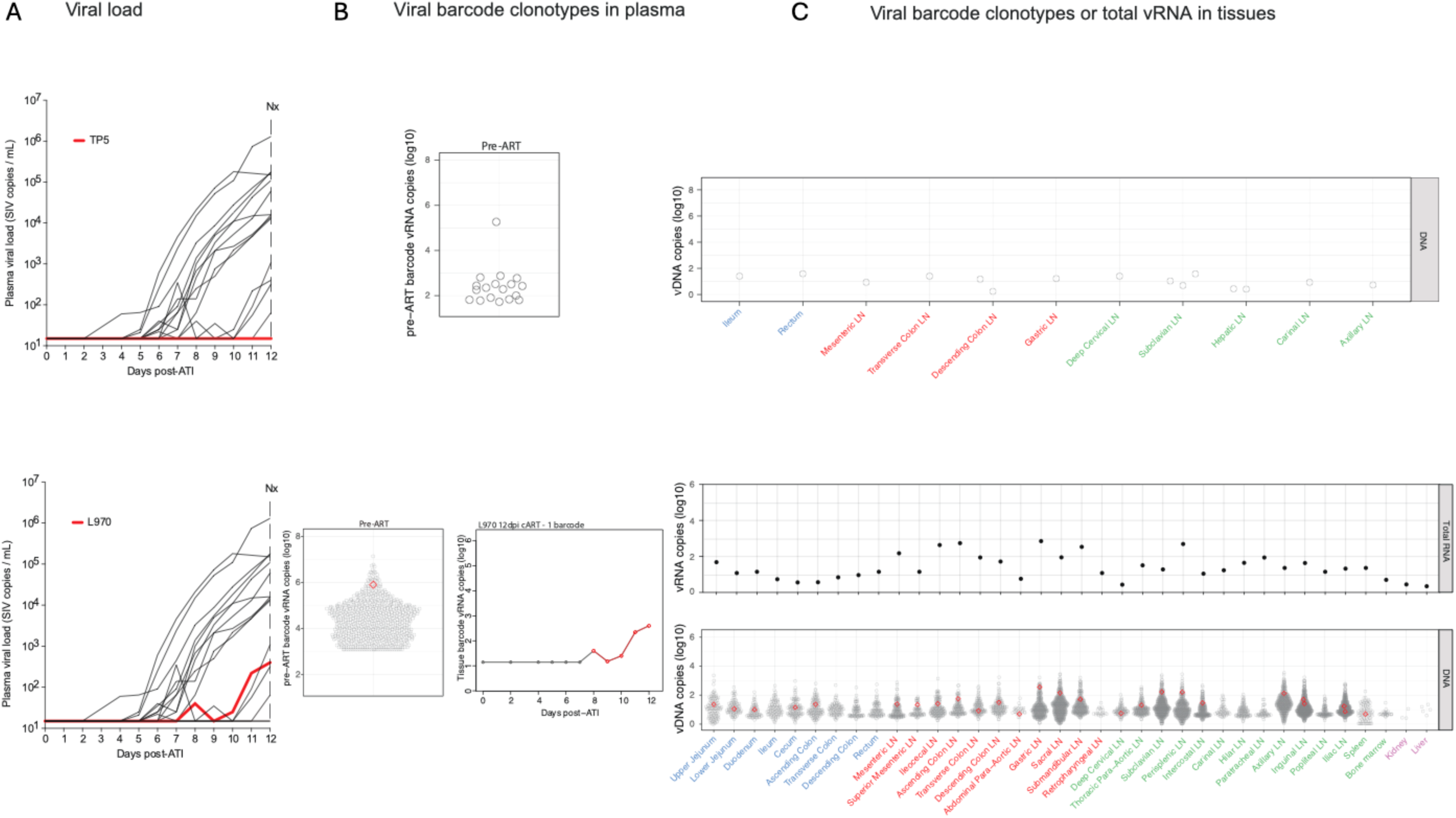
Viral barcode clonotypes in necropsy tissues from a non-rebounding RM TP5 and a single-barcode rebounder RM L970. (A) Plasma viral load dynamics following ART discontinuation for RMs TP5 (top) and L970 (bottom); red lines indicate individual viral load trajectories. RM TP5 exhibited no viral rebound through the 12-day monitoring period. (B) Proportional distributions of viral barcode clonotypes in plasma viremia during primary infection prior to ART initiation for RMs TP5 (top) and L970 (bottom); for L970, a single plasma barcode (BC.640), first detectable at day 11 following ART discontinuation is highlighted in red (bottom left panel). The bottom right panel shows calculated rebound viral growth curves for each rebounding barcode lineage with estimated time to a single copy in rebound viremia indicated. Red line indicates the dominant rebounding lineage. (C) Necropsy tissue distribution of vRNA and vDNA from primary infection for each animal. For RM L970, technical issues precluded vRNA barcode analysis in tissues, and thus vRNA values represent total vRNA rather than barcode vRNA. Grouped tissue categories are: GI tract (blue), GI tract draining lymph nodes (red), non-GI lymph tissues (green), non-lymphoid tissue (purple).

**Figure S8.**
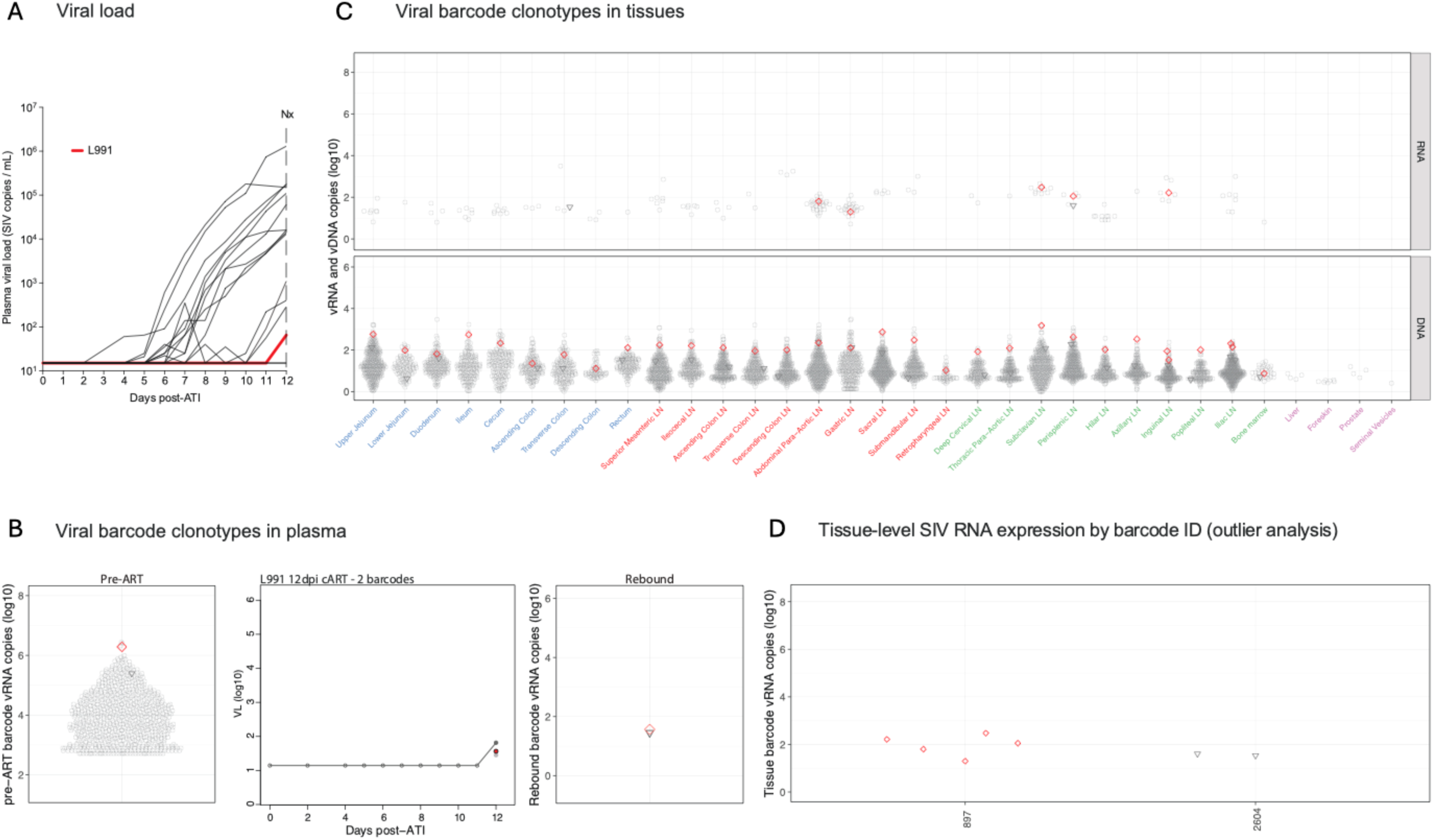
Viral barcode clonotypes in necropsy tissues and contributions to rebound viremia – RM L991, 2 rebounding barcodes. (A) Plasma viral load dynamics following ART discontinuation; red line indicates RM L991. (B) Left panel: Proportional distribution of viral barcode clonotypes in plasma viremia during primary infection prior to ART initiation, with the two barcodes found in rebound plasma at necropsy (BC.897 and BC.2604) highlighted. Middle panel: The calculated plasma viral load values for the two barcodes identified in the single day 12 timepoint, BC.897 and BC.2604, are shown. (C) Necropsy tissue distribution of vRNA and vDNA for the two barcodes identified in necropsy rebound plasma. Grouped tissue categories are: GI tract (blue), GI tract draining lymph nodes (red), non-GI lymph tissues (green), non-lymphoid tissue (purple). (D) Necropsy tissue vRNA levels for the two barcodes documented in rebound plasma viremia at necropsy; outlier analysis did not allow identification of presumptive tissue origin sites for either.

**Figure S9.**
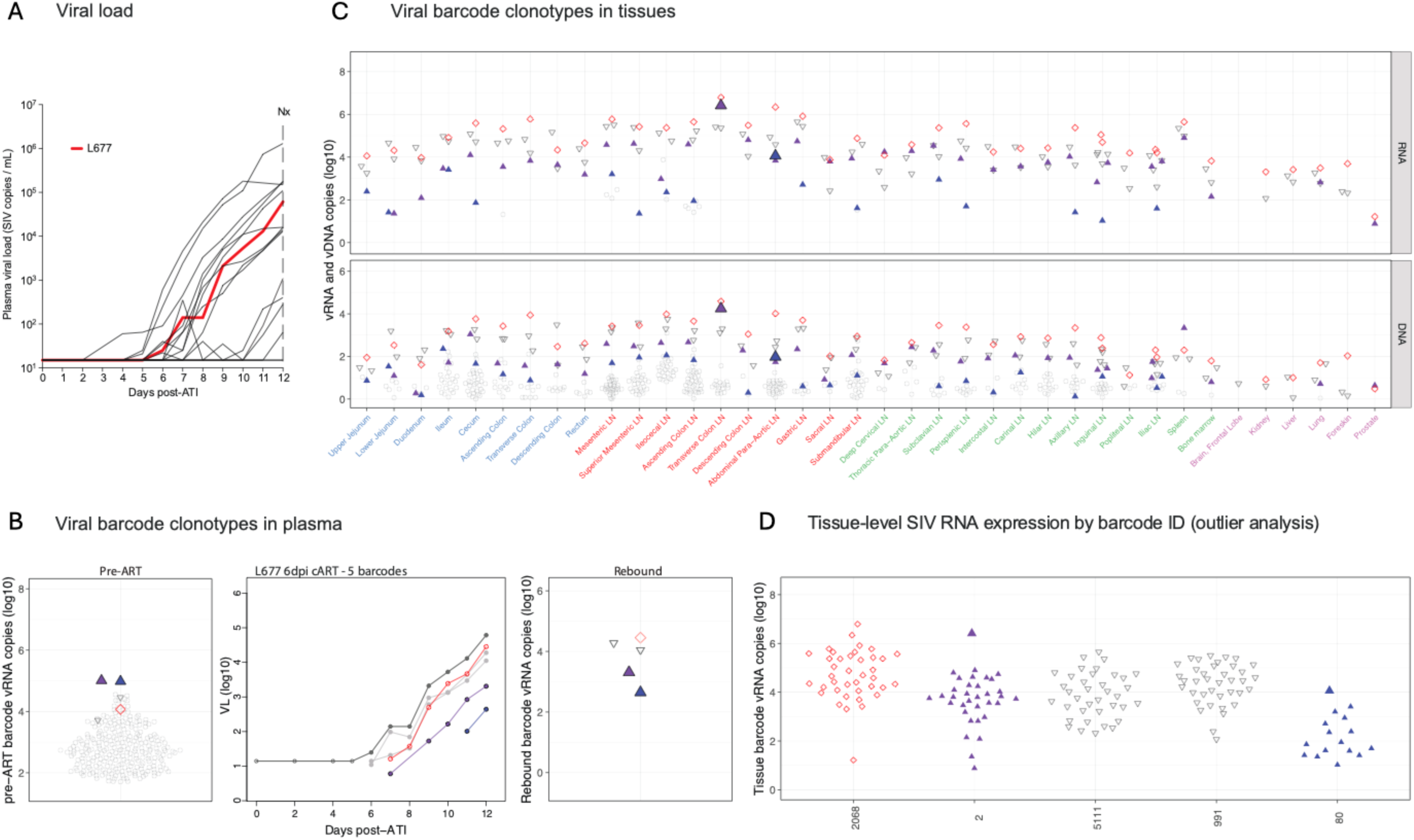
Viral barcode clonotypes in necropsy tissues and contributions to rebound viremia – RM L677, 5 rebounding barcodes. (A) Plasma viral load dynamics following ART discontinuation; red line indicates the viral load trajectory of RM L991. (B) Left panel: Proportional distribution of viral barcode clonotypes in plasma viremia during primary infection prior to ART initiation, with the 5 barcodes found in day 12 necropsy rebound plasma (BC.2068, BC.2, BC.5111, BC.001, and BC.80) highlighted. Middle panel: Calculated levels of plasma vRNA at necropsy for the 5 rebounding barcode lineage with estimated time to a single copy in rebound viremia indicated. Red line indicates the dominant rebounding lineage. Grey lines correspond to clones detected in rebound plasma but without an identified tissue origin site. Right panel: Proportional distribution of rebound viral barcode clonotypes in necropsy plasma at necropsy with barcodes for which a origin site was identified highlighted. (C) vRNA and vDNA distribution of the 5 rebounding barcodes (color-coded to panel B) in necropsy tissues. Grouped tissue categories are: GI tract (blue), GI tract draining lymph nodes (red), non-GI lymph tissues (green), non-lymphoid tissue (purple). Colored upward-facing triangles represent the 2 rebounding barcodes with tissue origin sites (BC.2 and BC.80) identified by outlier analysis indicated by the large, filled symbols. Unfilled symbols indicate barcodes for which outlier analysis did not identify a presumptive tissue origin site. (D) Viral barcode SIV RNA copies in tissue, grouped by barcode ID. Each point represents an individual barcode detected in necropsy tissue. Large, filled triangles indicate tissue barcode vRNA levels in tissue origin sites.

**Figure S10.**
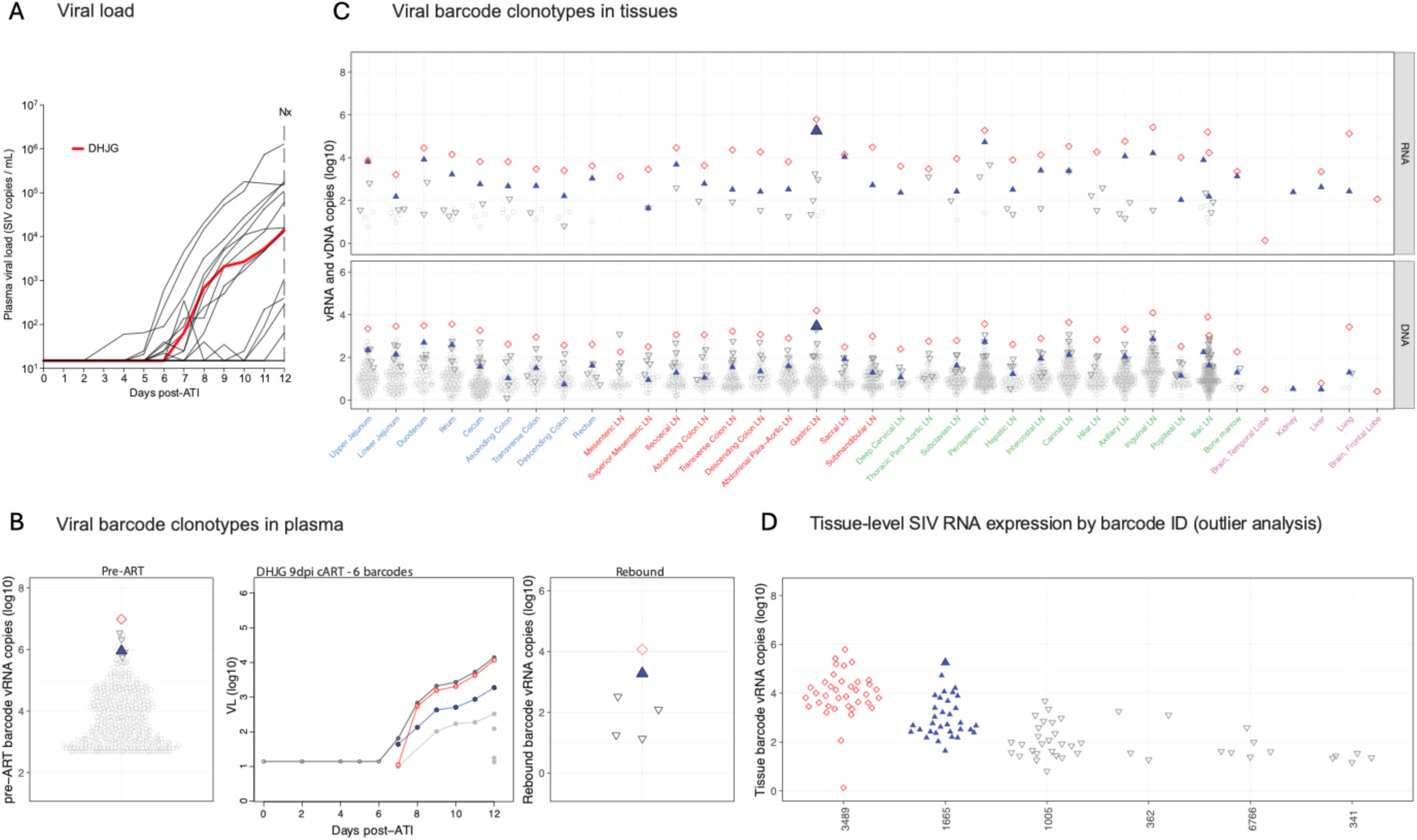
Viral barcode clonotypes in necropsy tissues and contributions to rebound viremia – RM DHJG, 6 rebounding barcodes. (A) Plasma viral load dynamics following ART discontinuation; red line indicates RM DHJG. (B) Left panel: Proportional distribution of viral barcode clonotypes in plasma viremia during primary infection prior to ART initiation, with barcodes found in rebound plasma highlighted. Middle panel: Calculated rebound viral growth curves for each rebounding barcode lineage with estimated time to a single copy in rebound viremia indicated. Red line indicates the dominant rebounding lineage (BC.3489). Grey lines correspond to barcodes detected in rebound plasma but without an identified presumptive tissue origin site. Right panel: Proportional distribution of rebound viral barcode clonotypes vRNA in necropsy plasma necropsy with one barcode with a tissue origin site identified by outlier analysis highlighted (large, filled triangle). (C) vRNA and vDNA distribution of all rebounding barcodes (color-coded to panel B) in necropsy tissues. Open symbols indicate barcodes without tissue origin site detected. Colored upward-facing triangles represent the one rebounding barcode with a tissue origin site in a gastric LN indicated by the large symbol. Grouped tissue categories are: GI tract (blue), GI tract draining lymph nodes (red), non-GI lymph tissues (green), non-lymphoid tissue (purple). (D) Viral barcode vRNA copies in necropsy tissue, grouped by barcode ID. Each point represents an individual barcode detected in tissue. Outlier analysis identified BC.1665 as the sole rebounding barcode for which a tissue origin site could be identified.

**Figure S11.**
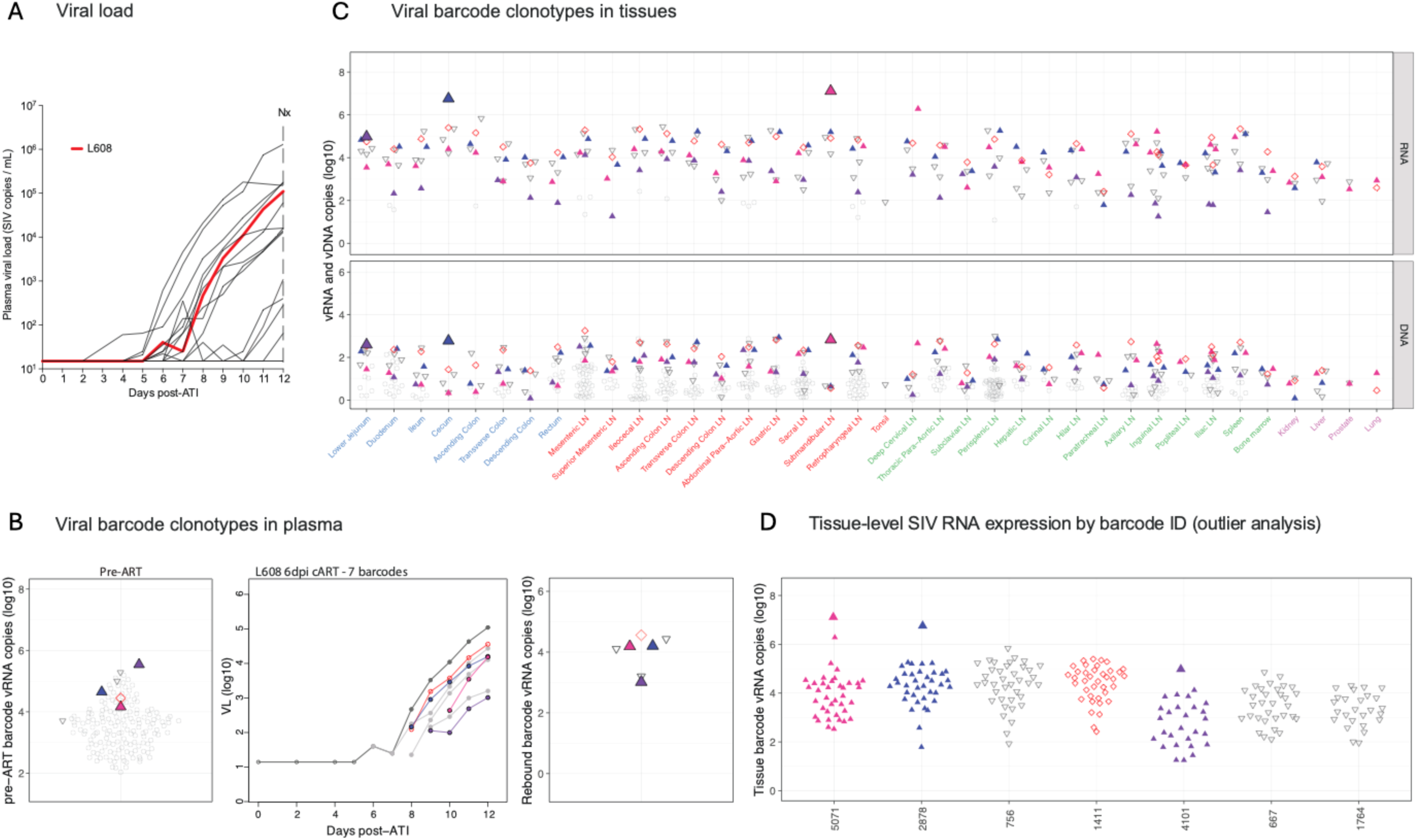
Viral barcode clonotypes in necropsy tissues and contributions to rebound viremia – RM L608, 7 rebounding barcodes. (A) Plasma viral load dynamics following ART discontinuation; red line indicates RM L608. (B) Left panel: Proportional distribution of all viral barcode clonotypes in plasma viremia during primary infection prior to ART initiation, with barcodes found in rebound plasma highlighted. Middle panel: Calculated rebound viral growth curves for each rebounding barcode lineage with estimated time to a single copy in rebound viremia indicated. Red line indicates the dominant rebounding lineage. Grey symbols correspond to barcodes detected in rebound plasma for which no presumptive tissue origin site was identified. Right panel: Proportional distribution of rebound plasma vRNA barcode clonotypes in necropsy plasma with barcodes for which tissue origin sites were identified highlighted. (C) vRNA and vDNA distribution of all rebounding barcodes (color-coded to panel B) in necropsy tissues. Colored upward-facing triangles represent rebounding barcodes with a tissue origin site, indicated by the large, filled triangles. Open symbols indicate barcodes without a presumptive tissue origin site identified. Grouped tissue categories are: GI tract (blue), GI tract draining lymph nodes (red), non-GI lymph tissues (green), non-lymphoid tissue (purple). (D) SIV RNA levels in necropsy tissues for barcodes present in rebound viremia, grouped by barcode ID. Each point represents an individual barcode detected in necropsy tissue. Tissue origin sites identified by outlier analysis for barcodes BC.5071, BC2.878, and BC.4101 are indicated by large, filled triangles.

**Figure S12.**
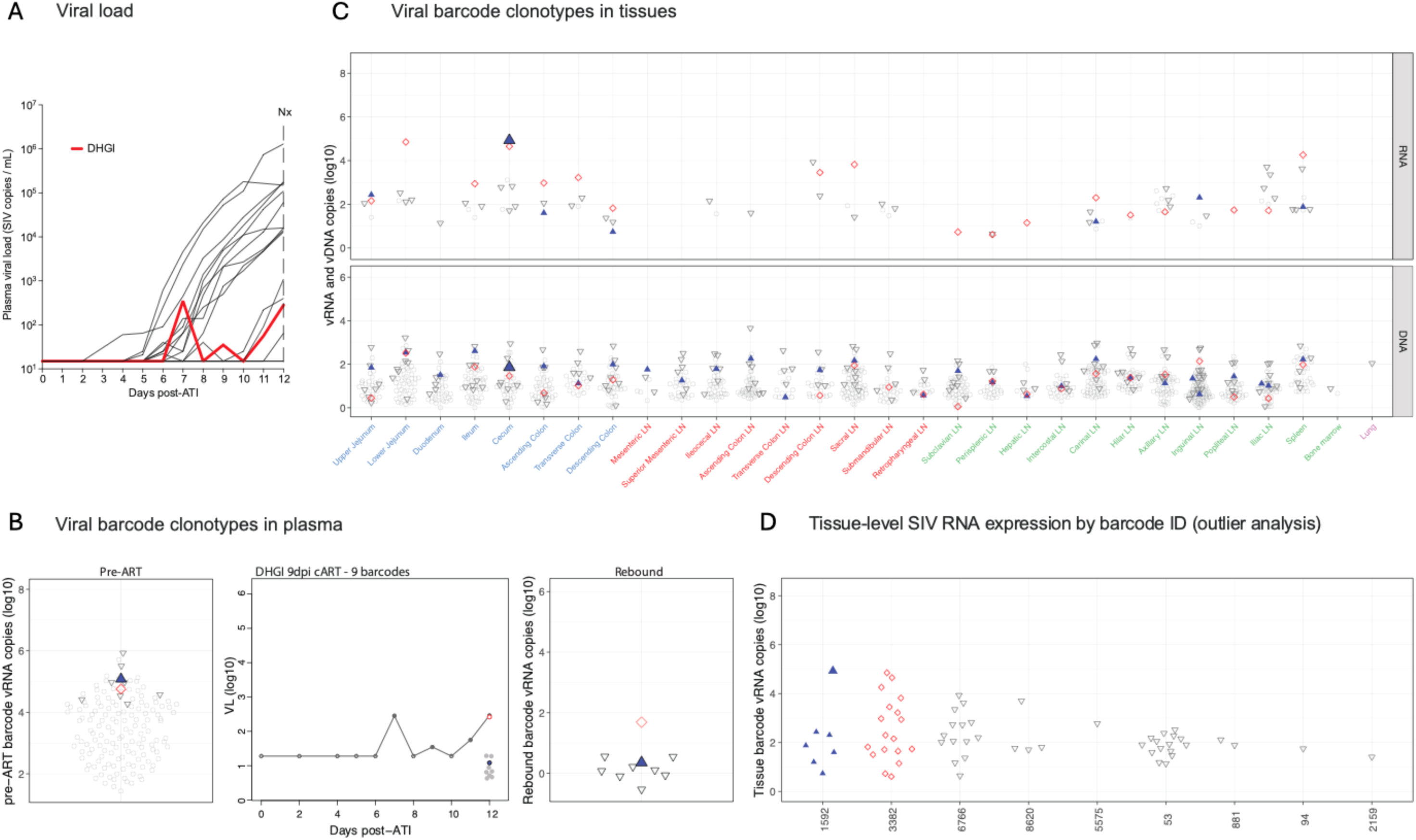
Viral barcode clonotypes in necropsy tissues and contributions to rebound viremia – RM DHGI, 9 rebounding barcodes. (A) Plasma viral load dynamics following ART discontinuation; red line indicates RM DHGI. (B) Left panel: Proportional distribution of viral barcode clonotypes in plasma viremia during primary infection prior to ART initiation, with all barcodes found in rebound plasma highlighted. Middle panel: Calculated rebound viral growth curves of each rebounding barcode lineage with estimated time to a single copy in rebound viremia indicated. Red line indicates the dominant rebounding lineage. Grey lines correspond to clones detected in rebound plasma but without an identified presumptive tissue origin site. Right panel: Proportional distribution of rebound viral barcode clonotypes in necropsy plasma with barcodes for which a tissue origin site was identified highlighted. (C) vRNA and vDNA distribution of all rebounding barcodes (color-coded to panel B) in necropsy tissues. Colored upward-facing triangles represent the one rebounding barcode (BC.1592) for which outlier analysis indicated a tissue origin site in cecum plotted with the by large, filled triangle. Open symbols indicate barcodes without presumptive tissue origin sites detected. Grouped tissue categories are: GI tract (blue), GI tract draining lymph nodes (red), non-GI lymph tissues (green), non-lymphoid tissue (purple). (D) Viral barcode SIV RNA copies in tissue, grouped by barcode ID. Each point represents an individual barcode detected in necropsy tissue, with the tissue origin for BC.1592 plotted as a filled triangle.

**Figure S13.**
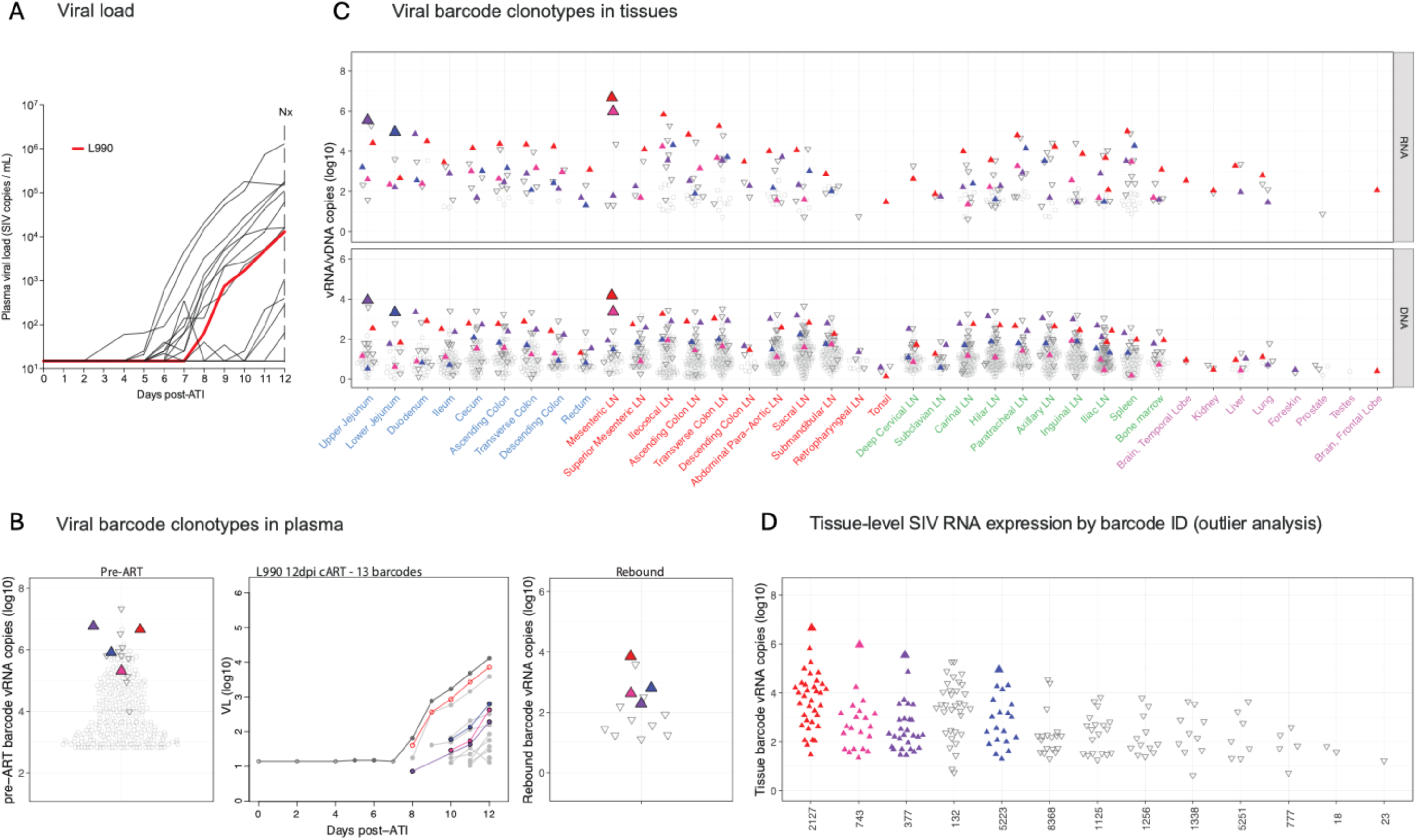
Viral barcode clonotypes in necropsy tissues and contributions to rebound viremia – RM L990, 13 rebounding barcodes. (A) Plasma viral load dynamics following ART discontinuation; red line indicates RM L990. (B) Left panel: Proportional distribution of viral barcode clonotypes in plasma viremia during primary infection prior to ART initiation, with all barcodes found in rebound plasma highlighted. Middle panel: Calculated rebound viral growth curves of each rebounding barcode lineage with estimated time to a single copy in rebound viremia indicated. Red line indicates the dominant rebounding lineage. Grey lineages correspond to barcodes detected in rebound plasma for which a presumptive tissue origin site was not identified. Right panel: Proportional distribution of rebound viral barcode clonotypes in plasma at necropsy with barcodes for which tissue origin sites were identified highlighted. (C) vRNA and vDNA distribution of all rebounding barcodes (color-coded to panel B) in necropsy tissues. Colored upward-facing triangles represent rebounding barcodes with tissue origin site indicated by the large symbol. Open symbols indicate barcodes without tissue origin site detected. Grouped tissue categories are: GI tract (blue), GI tract draining lymph nodes (red), non-GI lymph tissues (green), non-lymphoid tissue (purple). (D) Viral barcode SIV RNA copies in necropsy tissues, grouped by barcode ID. Each point represents an individual barcode detected in a necropsy tissue, with large, filled triangles indicating outliers allowing identification of tissue origin sites for BC.2127, BC.743, BC.377, and BC.5223.

**Figure S14.**
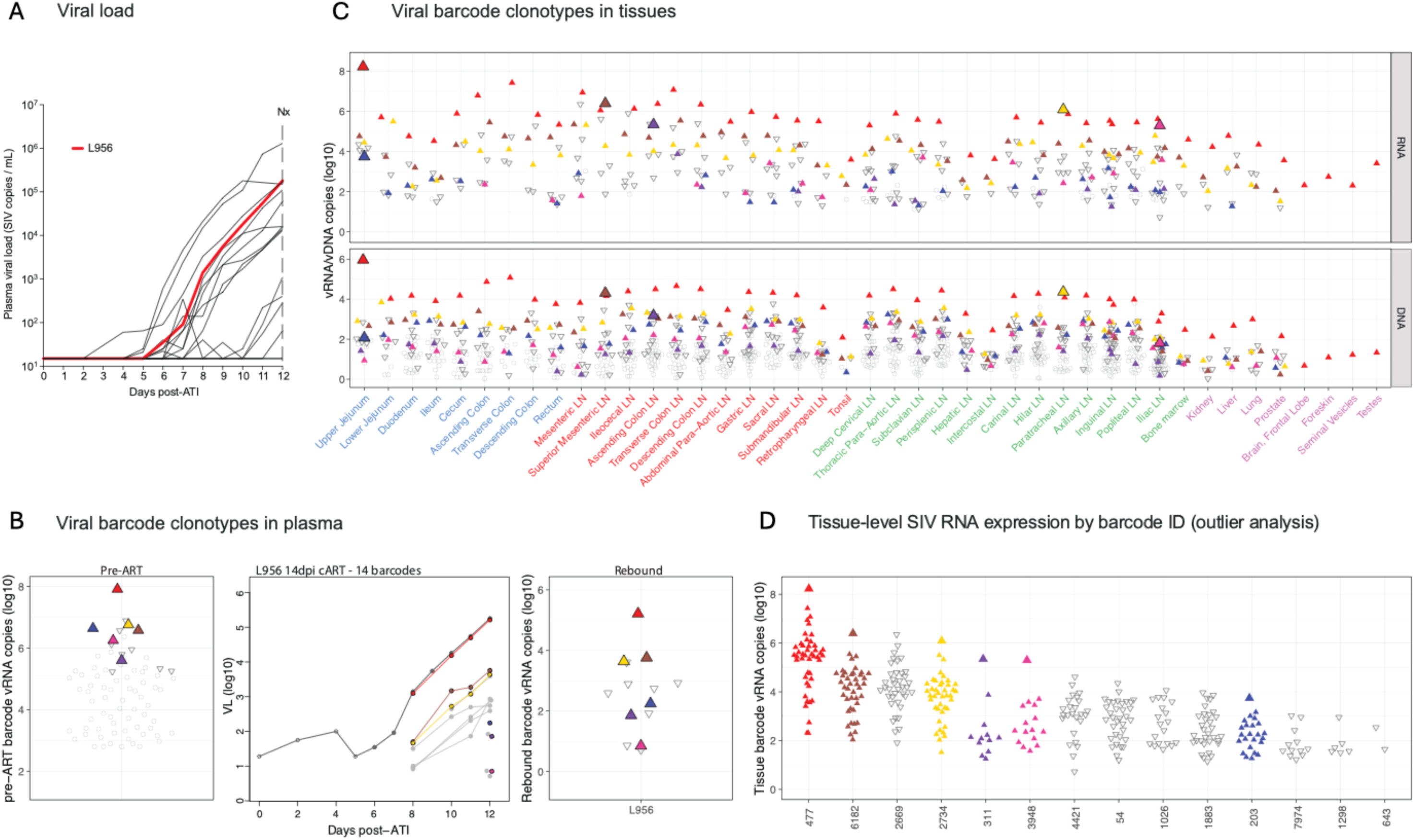
Viral barcode clonotypes in necropsy tissues and contributions to rebound viremia – RM L956, 14 rebounding barcodes. (A) Plasma viral load dynamics following ART discontinuation; red line indicates RM L956 (B) Left panel: Proportional distribution of viral barcode clonotypes in plasma viremia during primary infection prior to ART initiation, with all barcodes found in rebound plasma highlighted. Middle panel: Calculated rebound viral growth curves of each rebounding barcode lineage with estimated time to a single copy in rebound viremia indicated. Red line indicates the dominant rebounding lineage. Grey lines correspond to barcodes detected in rebound plasma for which a presumptive tissue origin site was not identified. Right panel: Proportional distribution of rebound viral barcode clonotypes in plasma at necropsy with barcodes having tissue origins highlighted. (C) vRNA and vDNA distribution of all rebounding barcodes (color-coded to panel B) in necropsy tissues. Colored upward-facing triangles represent rebounding barcodes with tissue origin sites indicated by large, filled triangles. Open symbols indicate barcodes without tissue origin site detected. Grouped tissue categories are: GI tract (blue), GI tract draining lymph nodes (red), non-GI lymph tissues (green), non-lymphoid tissue (purple). (D) Viral barcode SIV RNA copies in necropsy tissues, grouped by barcode ID. Each point represents an individual barcode detected in a necropsy tissue, with large, filled triangles indicating outliers allowing identification of tissue origin sites for BC.447, BC.6182, BC.2734, BC.311, BC.3948, and BC.203.

**Figure S15.**
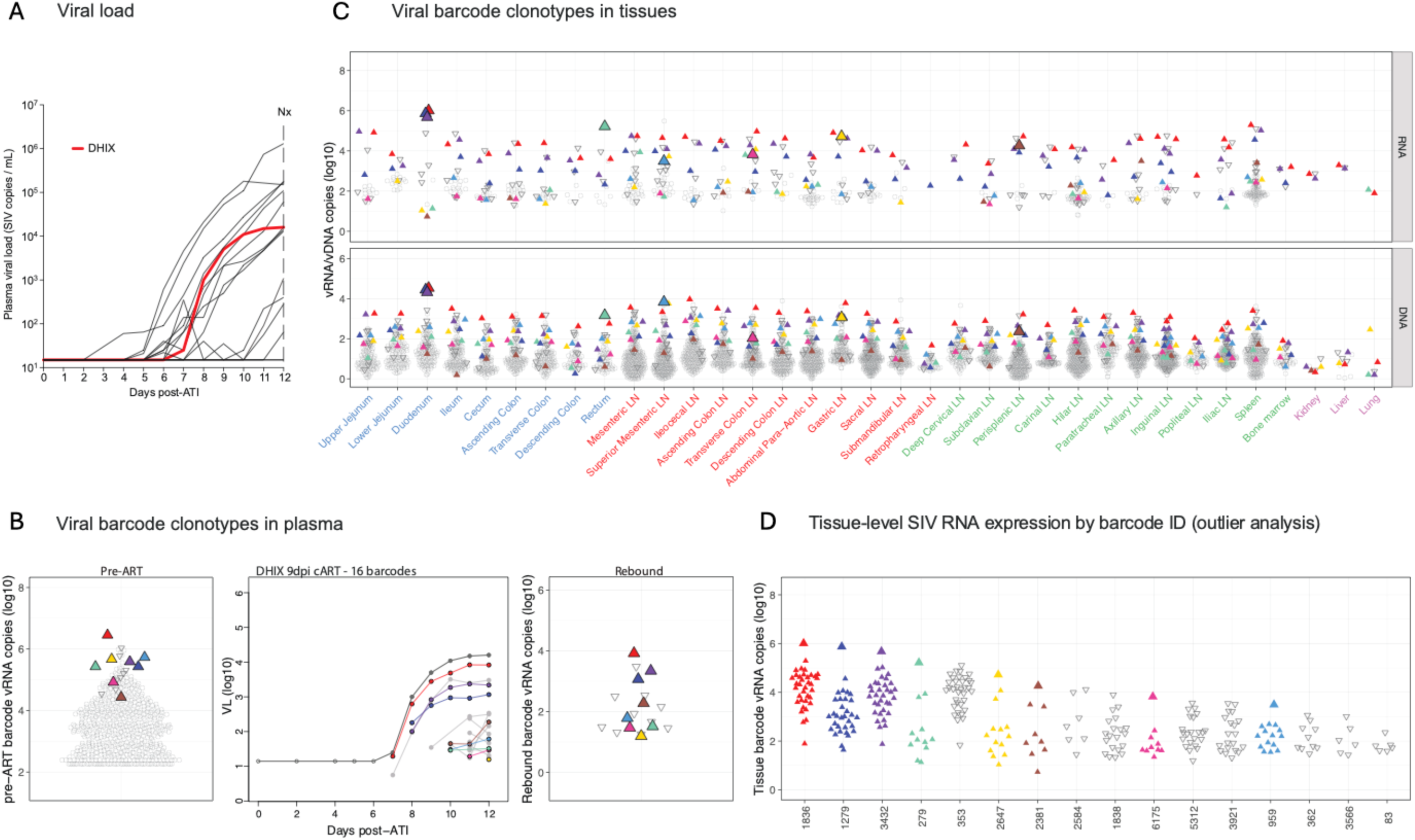
Viral barcode clonotypes in necropsy tissues and contributions to rebound viremia – RM DHIX, 16 rebounding barcodes. (A) Plasma viral load dynamics following ART discontinuation; red line indicates RM DHIX. (B) Left panel: Proportional distribution of viral barcode clonotypes in plasma viremia during primary infection prior to ART initiation, with all barcodes found in rebound plasma highlighted. Middle panel: Calculated rebound viral growth curves of each rebounding barcode lineage with estimated time to a single copy in rebound viremia indicated. Red line indicates the dominant rebounding lineage. Grey lines correspond to barcodes detected in rebound plasma for which a presumptive tissue origin site was not identified. Right panel: Proportional distribution of rebound viral barcode clonotypes in necropsy plasma with barcodes having tissue origin sites identified highlighted. (C) vRNA and vDNA distribution of all rebounding barcodes (color-coded to panel B) in necropsy tissues. Colored upward-facing triangles represent rebounding barcodes for which outlier analysis identified a tissue origin site, indicated by large, filled triangles. Open symbols indicate barcodes for which a tissue origin site was not detected. Grouped tissue categories are: GI tract (blue), GI tract draining lymph nodes (red), non-GI lymph tissues (green), non-lymphoid tissue (purple). (D) Viral barcode SIV RNA copies in necropsy tissues, grouped by barcode ID. Each point represents an individual barcode detected in a necropsy tissue. Large, filled triangles indicated tissue specimens identified by outlier analysis as tissue origin sites for barcodes BC.1836, BC.1279, BC.3432, BC.279, BC.2647, BC.2381, BC.6175, and BC.959 identified in necropsy rebound viremia.

**Figure S16.**
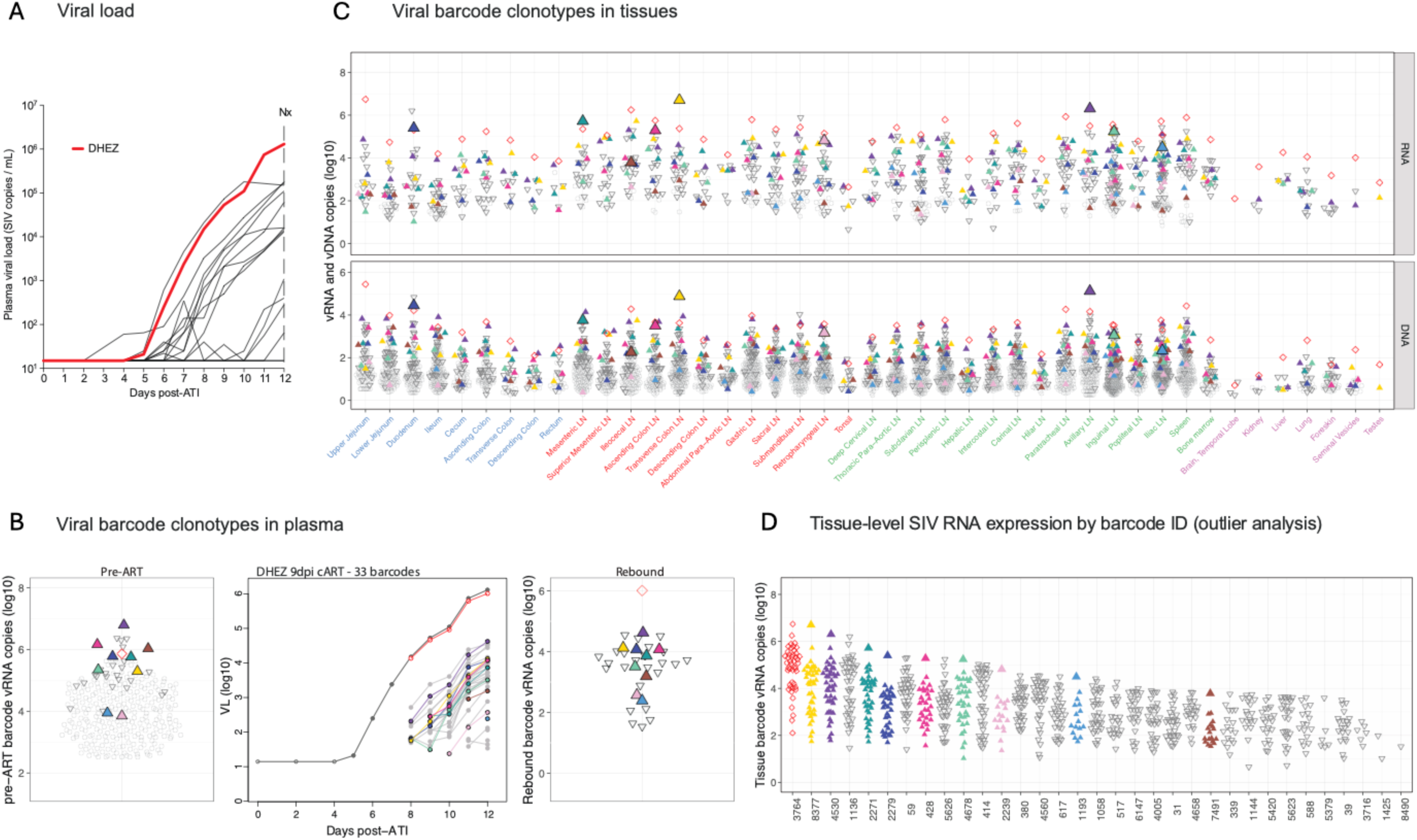
Viral barcode clonotypes in necropsy tissues and contributions to rebound viremia – RM DHEZ, 33 rebounding barcodes. (A) Plasma viral load dynamics following ART discontinuation; red line indicates RM DHEZ. (B) Left panel: Proportional distribution of viral barcode clonotypes in plasma viremia during primary infection prior to ART initiation, with all barcodes found in rebound plasma highlighted. Middle panel: Calculated rebound viral growth curves for each rebounding barcode lineage with estimated time to a single copy in rebound viremia indicated. Red line indicates the dominant rebounding lineage. Grey lines correspond to clones detected in rebound plasma for which no presumptive tissue origin site was identified. Right panel: Proportional distribution of rebound viral barcode clonotypes in necropsy plasma with barcodes for which a tissue origin site was identified highlighted. (C) vRNA and vDNA distribution of all rebounding barcodes (color-coded to panel B) in necropsy tissues. Colored upward-facing triangles represent rebounding barcodes with tissue origin site indicated by the large symbol. Open symbols indicate barcodes without tissue origin site detected. Grouped tissue categories are: GI tract (blue), GI tract draining lymph nodes (red), non-GI lymph tissues (green), non-lymphoid tissue (purple). (D) Viral barcode SIV RNA copies in necropsy tissues, grouped by barcode ID. Each point represents an individual barcode detected in a necropsy tissue. Large, filled triangles indicated tissue specimens identified by outlier analysis as tissue origin sites for barcodes BC.8377, BC.4530, BC.2271, BC.2279, BC.428, BC.4678, BC.2239, BC.1193, and BC.7491 identified in necropsy rebound viremia.

**Figure S17.**
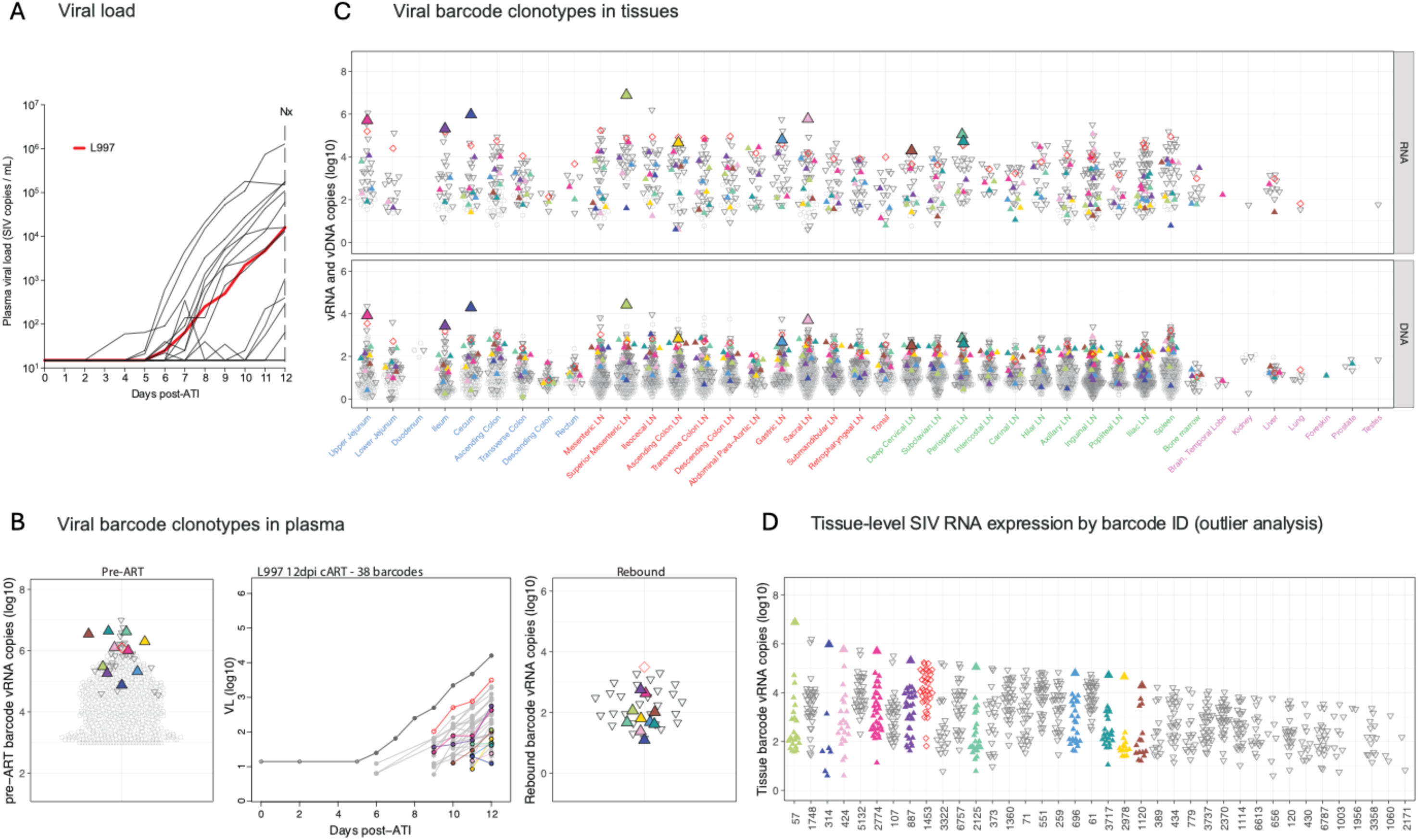
Viral barcode clonotypes in necropsy tissues and contributions to rebound viremia – RM L997, 38 rebounding barcodes. (A) Plasma viral load dynamics following ART discontinuation; red line indicates RM L997. (B) Left panel: Proportional distribution of viral barcode clonotypes in plasma viremia during primary infection prior to ART initiation, with all barcodes found in rebound plasma highlighted. Middle panel: Calculated rebound viral growth curves for each rebounding barcode lineage with estimated time to a single copy in rebound viremia indicated. Red line indicates the dominant rebounding lineage. Grey lines correspond to clones detected in rebound plasma for which no presumptive tissue origin site was identified. Right panel: Proportional distribution of rebound viral barcode clonotypes in necropsy plasma with barcodes for which a tissue origin site was identified highlighted. (C) vRNA and vDNA distribution of all rebounding barcodes (color-coded to panel B) in necropsy tissues. Colored upward-facing triangles represent rebounding barcodes with tissue origin site indicated by the large symbol. Open symbols indicate barcodes without tissue origin site detected. Grouped tissue categories are: GI tract (blue), GI tract draining lymph nodes (red), non-GI lymph tissues (green), non-lymphoid tissue (purple). (D) Viral barcode SIV RNA copies in necropsy tissues, grouped by barcode ID. Each point represents an individual barcode detected in a necropsy tissues, with large, filled triangles indicating tissue specimens identified in outlier analysis as tissue origins for barcodes contributing to rebound viremia, including BC.57, BC.314, BC.424, BC.2774, BC.887, BC.2125, BC.696, BC.3717, BC.2978, and BC.1120.

**Figure S18.**
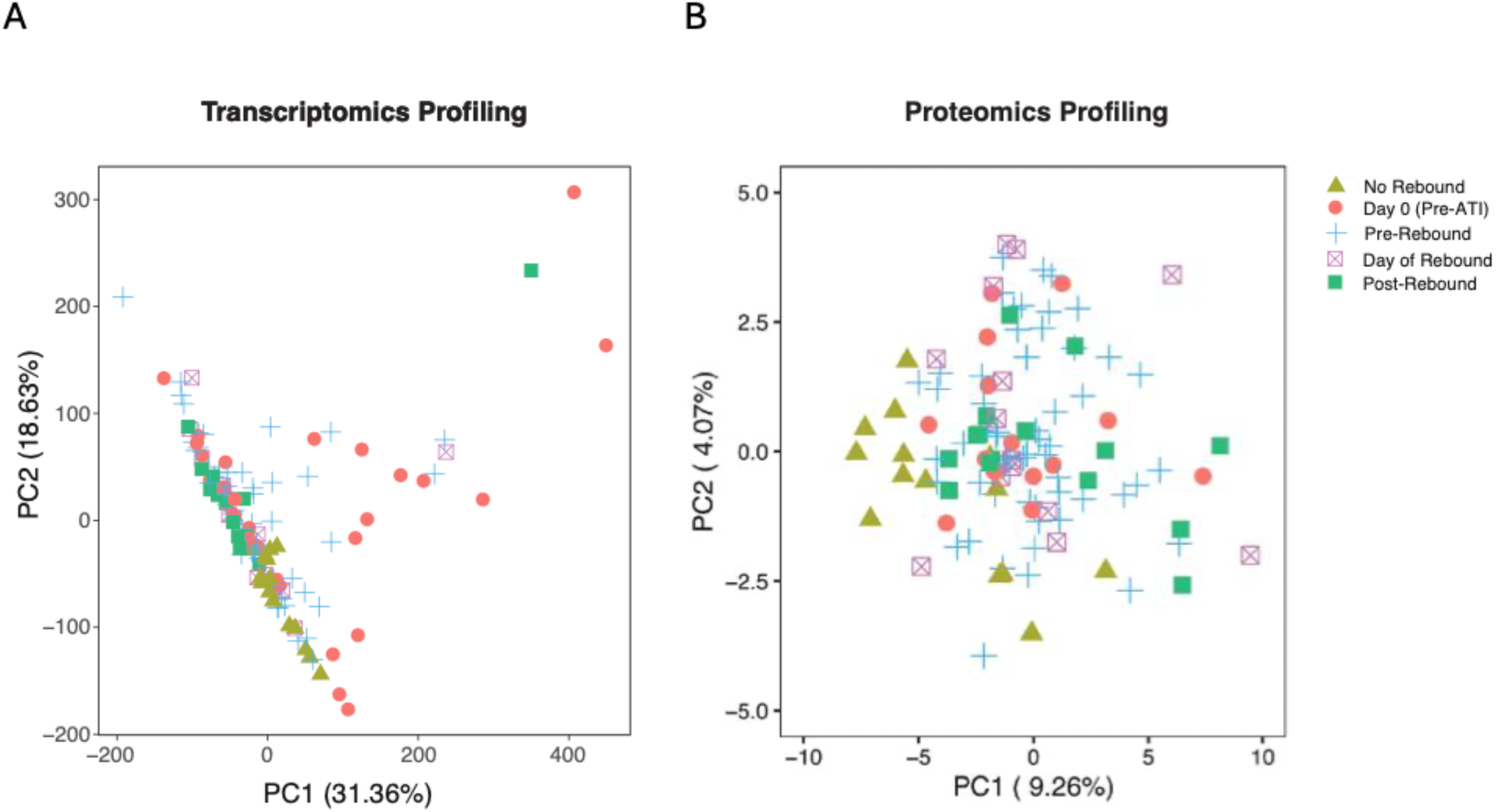
Principal component analysis of transcriptomic and proteomic data. Colors represent different timepoints following ART discontinuation relative to the day of rebound, defined for the purpose of this analysis as viral load >50 copies/mL.

**Figure S19.**
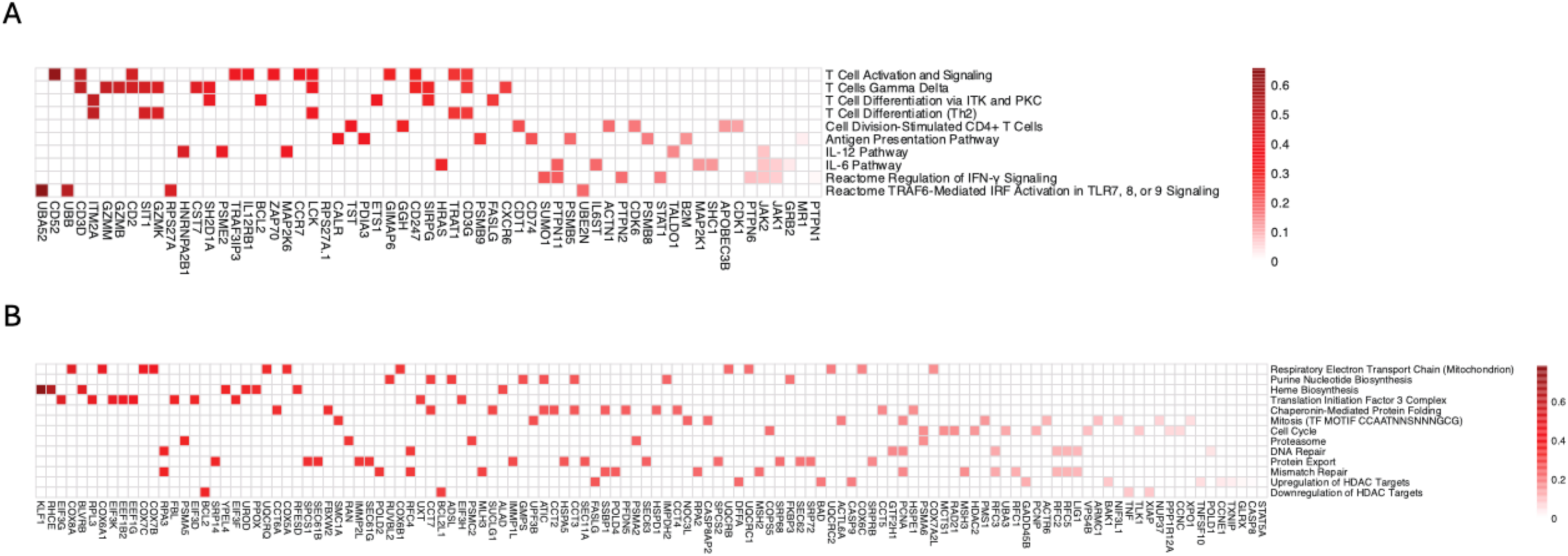
Genes driving transcriptomic upregulation in pathways at pre-rebound timepoints relative to baseline following ART discontinuation. Genes contributing to transcriptomic pathway upregulation in pre-rebound samples (viral load <50 cp/mL) compared to baseline (last timepoint prior to ATI) following ART discontinuation. (A) Genes driving upregulation of immune-related pathways and (B) genes associated with metabolism, cell cycle, and chromatin-remodeling pathways.

**Figure S20.**
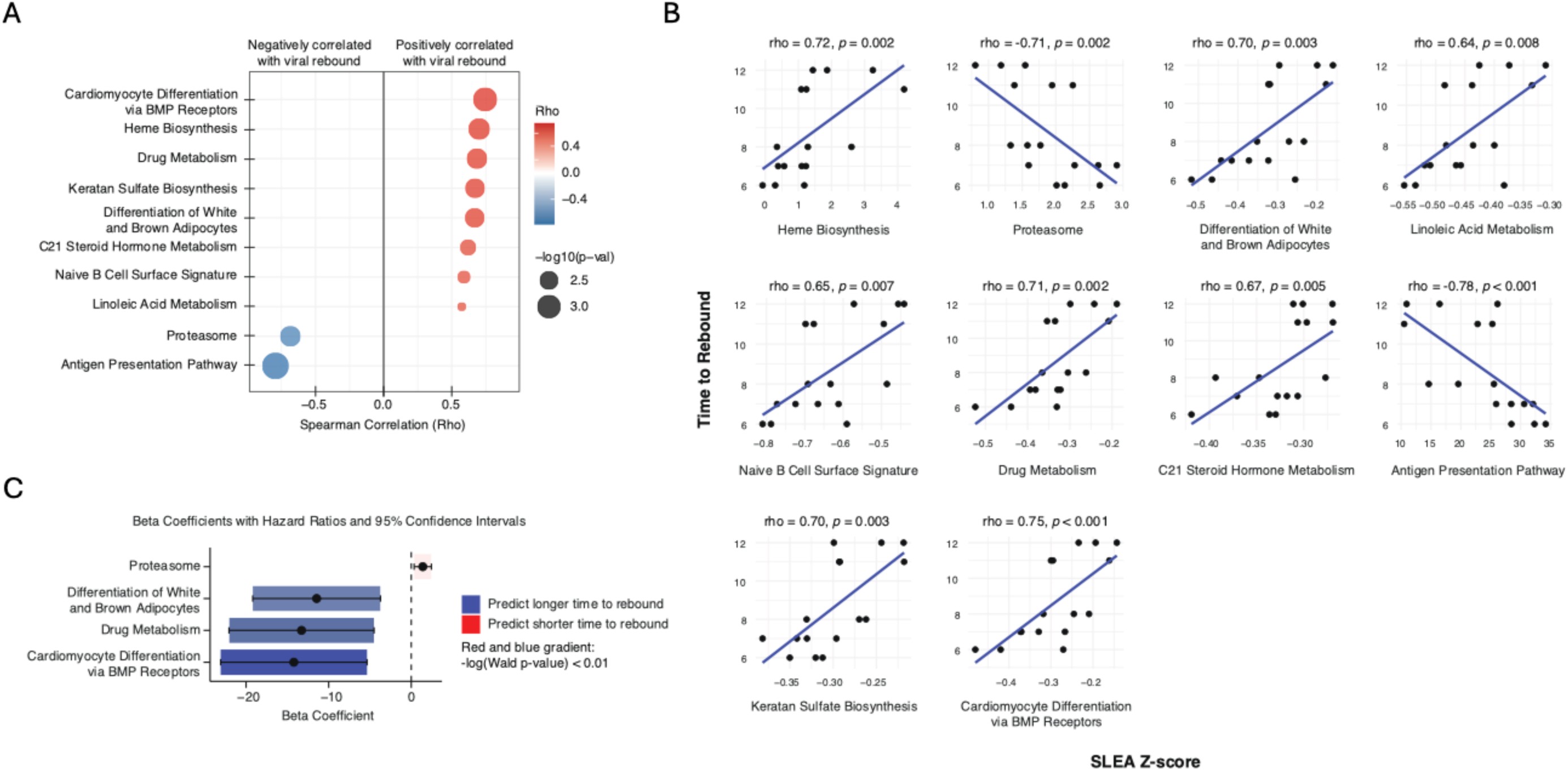
Correlation of transcriptomic pathways with time to viral rebound. (A) Spearman correlation analysis of transcriptomic pathways with time to viral rebound following ART discontinuation. Pathways with significant correlations (*p*<0.01) are shown, with red indicating positive correlations and blue indicating negative correlations. (B) Scatterplots of transcriptomic pathways correlated with time to viral rebound. The x-axis shows the SLEA Z-score at day 0, and the y-axis shows the time to viral rebound. Spearman correlations and *p*-values are shown for each pathway. (C) Univariate Cox proportional hazards model identifies pathways associated with shorter or longer times to viral rebound, displaying the beta coefficient and 95% confidence intervals. Red denotes pathways linked to shorter rebound times, while blue represents pathways associated with longer rebound times. Color gradient intensity reflects the significant level as Wald *p*<0.01.

**Figure S21.**
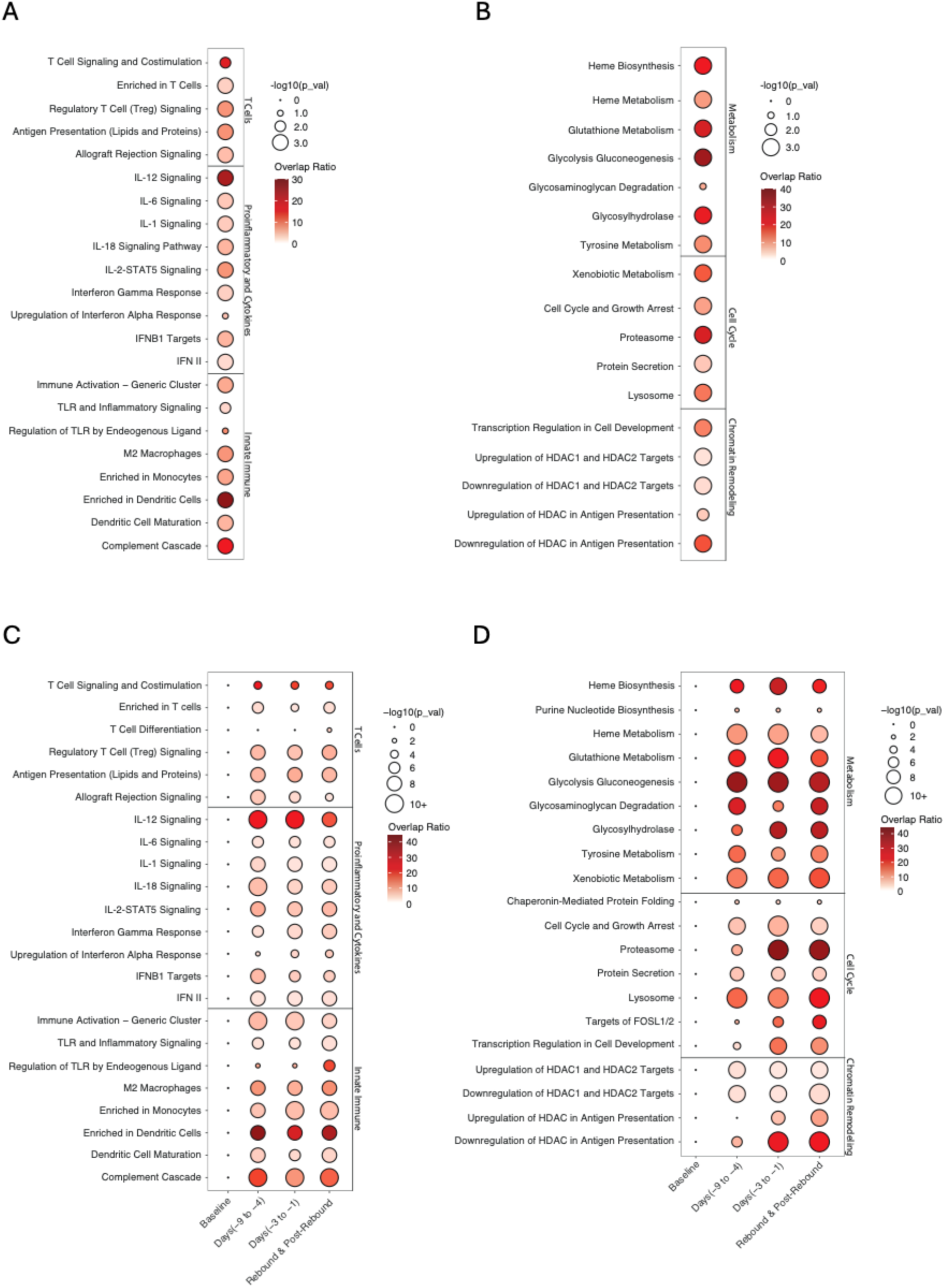
Proteomic changes following ART discontinuation before and after viral rebound relative to baseline. Upregulated proteomic pathways compared to baseline (last timepoint prior to ATI) are shown for pre-rebound samples (plasma viral loads <50 cp/mL) in (A) immune-related pathways and (B) metabolism, cell cycle, and chromatin remodeling pathways. Time course analysis depicts upregulated pathways relative to baseline for (C) immune pathways and (D) metabolism, cell cycle, and chromatin remodeling pathways across specific timepoints: baseline, days -9 to -4, days -3 to -1, and on the day of rebound and post-rebound >50 copies/mL. Each day is relative to the first timepoint with plasma viral load >50 copies/mL. Circle size reflects significance, while the color gradient represents the overlap ratio.

## Supplementary Table Legends

**Table S1.** Reactivation rates and viral dynamics across ART groups. Reactivation rates and viral rebound characteristics for SIV-infected macaques following ART discontinuation, organized by ART initiation group (day 6, day 9, day 12). Plasma viral load at necropsy (SIV copies/mL), rebound growth rate (log_10_ RNA/day), number of barcodes detected in plasma following ART discontinuation, and estimated reactivation rate (events/day) are reported for each animal. “N/A” indicates values not applicable for two animals (TP5 and DHHN) that did not rebound by the definition of viremia >50 copies/mL by day 12 following ART discontinuation.

| ART Group | Animal ID | Plasma Viral Load<br>at Necropsy<br>(SIV copies/mL) | Rebound Growth<br>Rate (log RNA/day) | Number of<br>Barcodes | Reactivation Rate<br>(events/day) |
| --- | --- | --- | --- | --- | --- |
| Day 6 | J639 | 1,100 | 2.03 | 1 | 0.12 |
|  | L604 | 180,000 | 2.22 | 3 | 0.49 |
|  | L608 | 110,000 | 1.58 | 7 | 0.84 |
|  | L677 | 61,000 | 1.29 | 5 | 0.6 |
|  | TP5 | < 50 | N/A | N/A | N/A |
| Day 9 | DHGI | 575 | 1.7 | 11 | 0.59 |
|  | DHEZ | 1,300,000 | 2.05 | 34 | 5.67 |
|  | DHIX | 16,000 | 2.65 | 8 | 1.38 |
|  | DHJG | 14,000 | 1.74 | 6 | 0.6 |
|  | DHHN | < 50 | N/A | N/A | N/A |
| Day 12 | L681 | 170,000 | 1.8 | 22 | 2.65 |
|  | L956 | 180,000 | 1.79 | 14 | 1.29 |
|  | L970 | 400 | 1.39 | 1 | 0.13 |
|  | L990 | 13,000 | 1.63 | 14 | 1.53 |
|  | L991 | 65 | 1.54 | 2 | 0.23 |
|  | L997 | 16,000 | 1.1 | 38 | 4.23 |

**Table S2.** RM demographics and tissues. Clinical data and the number of tissues collected per animal at necropsy.

| **Animal ID** | **Sex** | **Age at Study Start (Years)** | **Day of ART Initiation** | **Number of Tissues Collected at Necropsy** |
| --- | --- | --- | --- | --- |
| **DHEZ** | Male | 2.5 | 9 | 57 |
| **DHGI** | Male | 2.7 | 9 | 51 |
| **DHHN** | Male | 2.7 | 9 | 85 |
| **DHIX** | Male | 2.4 | 9 | 56 |
| **DHJG** | Male | 2.7 | 9 | 81 |
| **J639** | Male | 4.3 | 6 | 60 |
| **TP5** | Female | 4.5 | 6 | 54 |
| **L604** | Male | 2.5 | 6 | 62 |
| **L608** | Male | 2.5 | 6 | 71 |
| **L677** | Male | 2.4 | 6 | 76 |
| **L681** | Male | 2.5 | 12 | 52 |
| **L956** | Male | 2.4 | 12 | 62 |
| **L970** | Male | 2.4 | 12 | 73 |
| **L990** | Male | 2.4 | 12 | 55 |
| **L991** | Male | 2.5 | 12 | 62 |
| **L997** | Male | 2.4 | 12 | 71 |

**Table S3.**
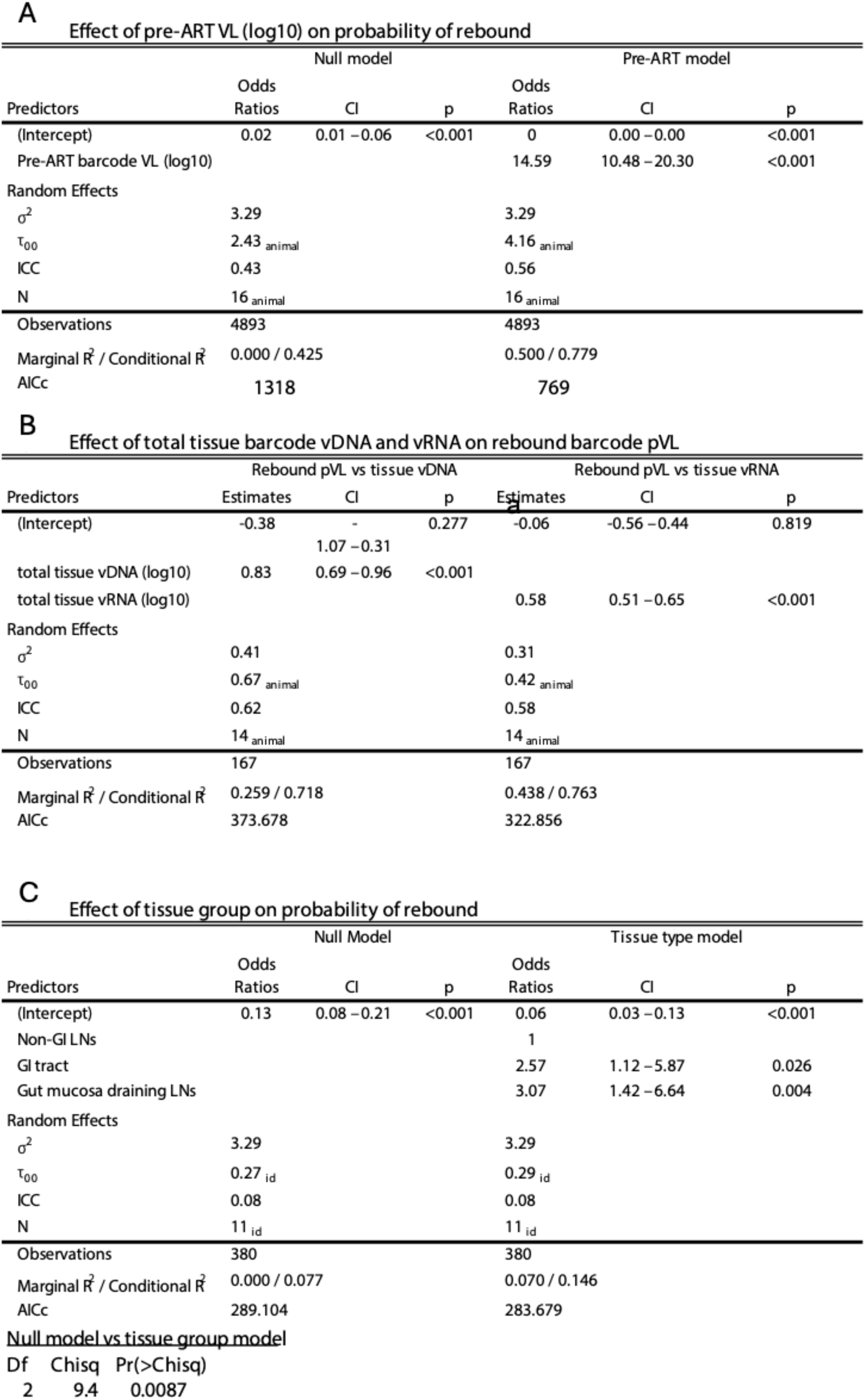
Linear regression and logistic regression models assessing virologic and tissue correlates of viral rebound. These tables summarize statistical models evaluating predictors of viral rebound following ART discontinuation in SIV-infected RMs. (A) Effect of barcode plasma viral loads prior to ART initiation on the probability of rebound using logistic regression. (B) Linear regression models relating total tissue-associated vDNA and vRNA levels (log_10_) to rebound barcode plasma viral loads. (C) Effect of anatomical tissue group (GI tract, GI-draining LNs, and non-GI LNs) on the probability of rebound using logistic regression. Odds ratios, 95% confidence intervals, and *p*-values are reported. Model fit parameters including AICc, variance components, and R^2^ values are included where applicable.

**Effect of pre-ART VL (log10) on probability of rebound**
| Predictors | Null model |  |  | Pre-ART model |  |  |
| --- | --- | --- | --- | --- | --- | --- |
|  | Odds Ratios | CI | p | Odds Ratios | CI | p |
| (Intercept) | 0.02 | 0.01 – 0.06 | <0.001 | 0 | 0.00 – 0.00 | <0.001 |
| Pre-ART barcode VL (log10) |  |  |  | 14.59 | 10.48 – 20.30 | <0.001 |
| Random Effects |  |  |  |  |  |  |
| $\sigma^2$ | 3.29 | | | 3.29 | | |
| $\tau_{00}$ | 2.43<sub>animal</sub> | | | 4.16<sub>animal</sub> | | |
| ICC | 0.43 |  |  | 0.56 |  |  |
| N | 16<sub>animal</sub> |  |  | 16<sub>animal</sub> |  |  |
| Observations | 4893 |  |  | 4893 |  |  |
| Marginal $R^2$ / Conditional $R^2$ | 0.000 / 0.425 | | | 0.500 / 0.779 | | |
| AICc | 1318 |  |  | 769 |  |  |

**B** **Effect of total tissue barcode vDNA and vRNA on rebound barcode pVL**
| Predictors | Rebound pVL vs tissue vDNA |  |  | Rebound pVL vs tissue vRNA |  |  |
| --- | --- | --- | --- | --- | --- | --- |
|  | Estimates | CI | p | Estimates | CI | p |
| (Intercept) | -0.38 | - | 0.277 | -0.06 | -0.56 – 0.44 | 0.819 |
|  |  | 1.07 – 0.31 |  |  |  |  |
| total tissue vDNA (log10) | 0.83 | 0.69 – 0.96 | <0.001 |  |  |  |
| total tissue vRNA (log10) |  |  |  | 0.58 | 0.51 – 0.65 | <0.001 |
| Random Effects |  |  |  |  |  |  |
| $\sigma^2$ | 0.41 | | | 0.31 | | |
| $\tau_{00}$ | 0.67<sub>animal</sub> | | | 0.42<sub>animal</sub> | | |
| ICC | 0.62 |  |  | 0.58 |  |  |
| N | 14<sub>animal</sub> |  |  | 14<sub>animal</sub> |  |  |
| Observations | 167 |  |  | 167 |  |  |
| Marginal $R^2$ / Conditional $R^2$ | 0.259 / 0.718 | | | 0.438 / 0.763 | | |
| AICc | 373.678 |  |  | 322.856 |  |  |

**C** **Effect of tissue group on probability of rebound**
| Predictors | Null Model |  |  | Tissue type model |  |  |
| --- | --- | --- | --- | --- | --- | --- |
|  | Odds Ratios | CI | p | Odds Ratios | CI | p |
| (Intercept) | 0.13 | 0.08 – 0.21 | <0.001 | 0.06 | 0.03 – 0.13 | <0.001 |
| Non-GI LNs |  |  |  | 1 |  |  |
| GI tract |  |  |  | 2.57 | 1.12 – 5.87 | 0.026 |
| Gut mucosa draining LNs |  |  |  | 3.07 | 1.42 – 6.64 | 0.004 |
| Random Effects |  |  |  |  |  |  |
| $\sigma^2$ | 3.29 | | | 3.29 | | |
| $\tau_{00}$ | 0.27<sub>id</sub> | | | 0.29<sub>id</sub> | | |
| ICC | 0.08 |  |  | 0.08 |  |  |
| N | 11<sub>id</sub> |  |  | 11<sub>id</sub> |  |  |
| Observations | 380 |  |  | 380 |  |  |
| Marginal $R^2$ / Conditional $R^2$ | 0.000 / 0.077 | | | 0.070 / 0.146 | | |
| AICc | 289.104 |  |  | 283.679 |  |  |

**Table S3.**
|  |  |  |
| --- | --- | --- |
| Df | Chisq | Pr(>Chisq) |
| 2 | 9.4 | 0.0087 |

## References

1. Ferris, A. L., Wells, D. W., Guo, S., Del Prete, G. Q., Swanstrom, A. E., Coffin, J. M., Wu, X., Lifson, J. D., and Hughes, S. H. (2019). Clonal expansion of SIV-infected cells in macaques on antiretroviral therapy is similar to that of HIV-infected cells in humans. PLoS Pathog. 15, e1007869. 10.1371/journal.ppat.1007869.

2. Okoye, A. A., Hansen, S. G., Vaidya, M., Fukazawa, Y., Park, H., Duell, D. M., Lum, R., Hughes, C. M., Ventura, A. B., Ainslie, E. et al. (2018). Early antiretroviral therapy limits SIV reservoir establishment to delay or prevent post-treatment viral rebound. Nature Medicine 24, 1430–1440. 10.1038/s41591-018-0130-7.

3. Chen, J., Zhou, T., Zhang, Y., Luo, S., Chen, H., Chen, D., Li, C., and Li, W. (2022). The reservoir of latent HIV. Front. Cell. Infect. Microbiol. 12, 1–15. 10.3389/fcimb.2022.945956.

4. Siliciano, J. D., and Siliciano, R. F. (2022). In Vivo Dynamics of the Latent Reservoir for HIV-1: New Insights and Implications for Cure. Annu. Rev. Pathol. Mech. Dis. 17, 271–294. 10.1146/annurev-pathol-050520-112001.

5. Pasternak, A. O., Grijsen, M. L., Wit, F. W., Bakker, M., Jurriaans, S., Prins, J. M., and Berkhout, B. (2020). Cell-associated HIV-1 RNA predicts viral rebound and disease progression after discontinuation of temporary early ART. JCI Insight 5, e134196. 10.1172/jci.insight.134196.

6. Fennessey, C. M., Pinkevych, M., Immonen, T. T., Reynaldi, A., Venturi, V., Nadella, P., Reid, C., Newman, L., Lipkey, L., Oswald, K. et al. (2017). Genetically-barcoded SIV facilitates enumeration of rebound variants and estimation of reactivation rates in nonhuman primates following interruption of suppressive antiretroviral therapy. PLoS Pathog. 13, e1006359. 10.1371/journal.ppat.1006359.

7. Liu, J., Ghneim, K., Sok, D., Bosche, W. J., Li, Y., Chipriano, E., Berkemeier, B., Oswald, K., Borducchi, E., Cabral, C. et al. (2016). Antibody-mediated protection against SHIV challenge includes systemic clearance of distal virus. Science 353, 6303. 10.1126/science.aag0491.

8. Del Prete, G. Q., Smedley, J., Macallister, R., Jones, G. S., Li, B., Hattersley, J., Zheng, J., Piatak Jr, M., Keele, B. F., Hesselgesser, J. et al. (2016). Short Communication: Comparative Evaluation of Coformulated Injectable Combination Antiretroviral Therapy Regimens in Simian Immunodeficiency Virus-Infected Rhesus Macaques. AIDS Research and Human Retroviruses 32, 163–168. 10.1089/AID.2015.0130.

9. Bruner, K. M., Wang, Z., Simonetti, F. R., Bender, A. M., Kwon, K. J., Sengupta, S., Fray, E. J., Beg, S. A., Antar, A. A. R., Jenike, K. M. et al. (2019). A novel quantitative approach for measuring the reservoir of latent HIV-1 proviruses. Nature 566, 120–125. 10.1038/s41586-019-0898-8.

10. Pace, M. J., Graf, E. H., Agosto, L. M., Mexas, A. M., Male, F., Brady, T., Bushman, F. D., and O’Doherty, U. (2012). Directly Infected Resting CD4+T Cells Can Produce HIV Gag without Spreading Infection in a Model of HIV Latency. PLoS Pathogens 8, e1002818. 10.1371/journal.ppat.1002818.

11. Keele, B. F., Okoye, A. A., Fennessey, C. M., Varco-Merth, B., Immonen, T. T., Kose, E., Conchas, A., Pinkevych, M., Lipkey, L., Newman, L. et al. (2024). Early antiretroviral therapy in SIV-infected rhesus macaques reveals a multiphasic, saturable dynamic accumulation of the rebound competent viral reservoir. PLoS Pathogens 20, e1012135. 10.1371/journal.ppat.1012135.

12. Picker, L., Keele, B., Okoye, A., Immonen, T., Varco-Merth, B., Duell, D., Nkoy, C., Goodwin, W., Hoffmeister, S., Kose, E. et al. (2025). Identification of initial sites of SIV rebound after treatment cessation. Preprint at Res Sq, 10.21203/rs.3.rs-6814218/v1.

13. Whitney, J. B., Lim, S., Osuna, C. E., Kublin, J. L., Chen, E., Yoon, G., Liu, P., Abbink, P., Borducci, E. N., Hill, A. et al. (2018). Prevention of SIVmac251 reservoir seeding in rhesus monkeys by early antiretroviral therapy. Nature Communications 9, 5429. 10.1038/s41467-018-07881-9.

14. Cadena, A. M., Ventura, J. D., Abbink, P., Borducchi, E. N., Tuyishime, H., Mercado, N. B., Walker-Sperling, V., Siamatu, M., Liu, P., Chandrashekar, A. et al. (2021). Persistence of viral RNA in lymph nodes in ART-suppressed SIV/SHIV-infected Rhesus Macaques. Nature Communications 12, 1471. 10.1038/s41467-021-21724-0.

15. Scheerder, M. D., Vrancken, B., Dellicour, S., Schlub, T., Lee, E., Shao, W., Rutsaert, S., Verhofstede, C., Kerre, T., Malfait, T. et al. (2019). HIV Rebound Is Predominantly Fueled by Genetically Identical Viral Expansions from Diverse Reservoirs. Cell Host & Microbe 26, 347–358. 10.1016/j.chom.2019.08.003.

16. Barouch, D. H., Ghneim, K., Bosche, W. J., Li, Y., Berkemeier, B., Hull, M., Bhattacharyya, S., Cameron, M., Liu, J., Smith, K. et al. (2016). Rapid Inflammasome Activation Following Mucosal SIV Infection of Rhesus Monkeys. Cell 165, 656–667. 10.1016/j.cell.2016.03.021.

17. Giron, L. B., Palmer, C. S., Liu, Q., Yin, X., Papasavvas, E., Sharaf, R., Etemad, B., Damra, M., Goldman, A. R., Tan, H. et al. (2021). Non-invasive plasma glycomic and metabolic biomarkers of post-treatment control of HIV. Nature Communications 12, 3922. 10.1038/s41467-021-24077-w.

18. Julg, B., Walker-Sperling, V. E. K., Wagh, K., Aid, M., Stephenson, K. E., Zash, R., Liu, J., Nkolola, J. P., Hoyt, A., Castro, M. et al. (2024). Safety and antiviral effect of a triple combination of HIV-1 broadly neutralizing antibodies: a phase 1/2a trial. Nature Medicine. 10.1038/s41591-024-03247-5.

19. Bolton, D. L., Pegu, A., Wang, K., McGinnis, K., Nason, M., Foulds, K., Letukas, V., Schmidt, S. D., Chen, X., Todd, J. P. et al. (2016). Human Immunodeficiency Virus Type 1 Monoclonal Antibodies Suppress Acute Simian-Human Immunodeficiency Virus Viremia and Limit Seeding of Cell-Associated Viral Reservoirs. Journal of Virology 90. 10.1128/JVI.02454-15.

20. Li, H., Wang, S., Kong, R., Ding, W., Lee, F., Parker, Z., Kim, E., Learn, G. H., Hahn, P., Policicchio, B. et al. (2016). Envelope residue 375 substitutions in simian-human immunodeficiency viruses enhance CD4 binding and replication in rhesus macaques. PNAS 113, e3413–e3422. 10.1073/pnas.1606636113.

21. Pinkevych, M., Fennessey, C. M., Cromer, D., Reid, C., Trubey, C. M., Lifson, J. D., Keele, B. F., and Davenport, M. P. (2019). Predictors of SIV recrudescence following antiretroviral treatment interruption. eLife 8, e49022. 10.7554/eLife.49022.

22. Hansen, S. G., Ford, J. C., Lewis, M. S., Ventura, A. B., Hughes, C. M., Coyne-Johnson, L., Whizin, N., Oswald, K., Shoemaker, R. Swanson, T., et al. (2011). Profound early control of highly pathogenic SIV by an effector memory T-cell vaccine. Nature 473, 523–527. 10.1038/nature10003.

23. Bender, A. M., Simonetti, F. R., Kumar, M. R., Fray, E. J., Bruner, K. M., Timmons, A. E., Tai, K. Y., Jenike, K. M., Antar, A. A. R., Liu, P. et al. (2019). The Landscape of Persistent Viral Genomes in ART-Treated SIV, SHIV, and HIV-2 Infections. Cell Host Microbe *26*, 73-85. 10.1016/j.chom.2019.06.005.

24. Gaebler, C., Falcinelli, S. D., Stoffel, E., Read, J., Murtagh, R., Oliveira, T. Y., Ramos, V., Lorenzi, J. C. C., Kirchherr, J., James, K. S. et al. (2021). Sequence Evaluation and Comparative Analysis of Novel Assays for Intact Proviral HIV-1 DNA. Journal of Virology 95. 10.1128/JVI.01986-20.

